# Evidence from combined analysis of single cell RNA-Seq and ATAC-Seq data of regulatory toggles operating in native and iPS-derived murine retina

**DOI:** 10.1101/2020.03.02.972497

**Authors:** Anouk Georges, Arnaud Lavergne, Michiko Mandai, Fanny Lepiemme, Latifa Karim, Loic Demeulenaere, Diego Aguilar, Michael Schyns, Sébastien Dupont, Laurent Nguyen, Jean-Marie Rakic, Masayo Takahashi, Michel Georges, Haruko Takeda

## Abstract

We report the generation and analysis of single-cell RNA-Seq data (> 38,000 cells) from native and iPSC-derived murine retina at four matched developmental stages spanning the emergence of the major retinal cell types. We combine information from temporal sampling, visualization of 3D UMAP manifolds, pseudo-time and RNA velocity analyses, to show that iPSC-derived 3D retinal aggregates broadly recapitulate the native developmental trajectories. However, we show relaxation of spatial and temporal transcriptome control, premature emergence and dominance of photoreceptor precursor cells, and susceptibility of dynamically regulated pathways and transcription factors to culture conditions in iPSC-derived retina. We generate bulk ATAC-Seq data for native and iPSC-derived murine retina identifying ∼125,000 peaks. We combine single-cell RNA-Seq with ATAC-Seq information and obtain evidence that approximately half the transcription factors that are dynamically regulated during retinal development may act as repressors rather than activators. We propose that sets of activators and repressors with cell-type specific expression constitute “regulatory toggles” that lock cells in distinct transcriptome states underlying differentiation. We provide evidence supporting our hypothesis from the analysis of publicly available single-cell ATAC-Seq data for adult mouse retina. We identify subtle but noteworthy differences in the operation of such toggles between native and iPSC-derived retina particularly for the Etv1, Etv5, Hes1 and Zbtb7a group of transcription factors.

## Introduction

It has recently become possible to recapitulate retinal development from induced pluripotent stem cells (iPSCs) in human and mice [1-3]. This has opened new avenues to explore the molecular mechanisms underlying developmental competence, commitment and differentiation for each of the major cell types during retinal neurogenesis. It offers hope to improve therapies for retinal degenerative diseases which afflict tens of millions of people in the US and Europe alone and may account for approximately 50% of all cases of blindness [4]. Stem cells derived from patient-specific somatic cells offer new opportunities to study the effects of gene defects on human retinal development in vitro and to test small molecules or biologics to treat the corresponding disorders [5,6].

Assessing how faithfully iPSC-derived 3D retinal aggregates recapitulate specific developmental programs has typically been done by monitoring the expression of limited numbers of cell-type specific markers and examining the spatial patterning of the corresponding groups of cells [7]. Interrogating the expression of a handful of marker genes/proteins does not fully inform about the proper temporal and spatial execution of the epigenetic program, nor does it inform about the presence of aberrant cell types. Single-cell RNA-sequencing (scRNA-Seq) now enables the profiling of samples of the transcriptome (typically between 3% and 15% of mRNAs present in a cell depending on the methodology) of individual cells. This permits the clustering of cells based on the similarity of their transcriptome and the identification of cellular subtypes including some that may not have been recognized before [8]. It allows to refine developmental trajectories by identifying cells occupying intermediate states connecting clusters in multidimensional expression space [9,10] and by predicting the developmental orientation taken by individual cells based on measured deviations from the steady-state ratio between spliced and unspliced RNA molecules (“RNA velocity”) [11,12]. Genes that are defining cellular sub-types can be pinpointed by differential expression analysis between clusters [13], while genes that drive the differentiation process may be identified by searching for gene sets that are dynamically regulated across real and/or pseudo-time [14]. Recently, scRNA-Seq has been used to compare transcriptome dynamics during native and embryonic stem cells (ESC)- or iPSC-derived retinal development in human [15-17]. This has revealed comparable cellular composition at equivalent ages and the convergence of the organoid transcriptomes to that of adult peripheral retinal cell types with, however, some differences in gene expression of particular cell types as well as structural differences of inner retinal lamination that seems disrupted in advanced organoid stages compared with fetal retina [16]. It has revealed striking cell type-specific expression of genes underpinning inherited diseases such as Leber congenital amaurosis, retinitis pigmentosa, stationary night blindness and achromatopsia, and its conservation in organoids [17].

Here we report the generation and use of scRNA-Seq data collected at four matched stages of native and iPSC-derived retinal development in the mouse to study the dynamics of the transcriptome and compare it between the two systems. We integrate scRNA-Seq data with bulk and single-cell ATAC-seq data (which identify active gene regulatory elements by virtue of local chromatin openness [18]), and provide evidence for the operation of transcription factor (TF)-based regulatory toggles combining activators and repressors, that may lock the transcriptome of distinct cellular sub-types in both native and iPSC-derived retina thereby underpinning the different cellular identities.

## Results

### Joint analysis of scRNA-Seq data from native retina and iPSC-derived 3D retinal aggregates highlights canonical cell types and developmental trajectories

To contribute to the comparison of the developmental trajectories in native retina (NaR) and iPSC-derived 3D retinal aggregates (3D-RA), we performed scRNA-Seq of murine NaR and 3D-RA at four matched stages of development: embryonic day (E)13 vs differentiation day (DD)13, postnatal day (P)0 vs DD21, P5 vs DD25 and P9 vs DD29 [19]. NaR were dissected from two to 11 C57BL/6 mice (of both sexes) per stage. Mouse 3D-RA were generated from the Nrl-GFP (C57BL/6 background) iPSC line [20] following [21-22] (SFig. 1). Optic vesicle-like structures (OV) were manually dissected from 3D-RA. Cells from NaR and OV were dissociated and subjected to droplet-based scRNA-Seq using a 10X Genomics Chromium platform. We obtained sequence information for 21,249 cells from NaR and 16,842 cells from 3D-RA, distributed evenly amongst developmental stages. We generated an average of 74,808 reads per cell, corresponding to 5,940 unique molecular identifiers (UMIs) and 2,471 genes per cell (STable 1).

We first analyzed all data jointly (i.e. NaR and 3D-RA) to cover a maximum of intermediate developmental stages and hence generate the most continuous manifold possible. We used Canonical Correlation Analysis (CCA) implemented with Seurat [23] to align the NaR and 3D-RA datasets based on the expression profiles of 1,253 “most variable” genes (STable 2). We projected the corresponding 30-dimensional distances (based on the 30 first CCA) between cells in 2D- and 3D-space using Uniform Manifold Approximation and Projection (UMAP) [24]. We assigned all 38,091 cells jointly (i.e. NaR and 3D-RA) to 71 clusters by k-means clustering (Fig. 1A).

**Figure 1:**
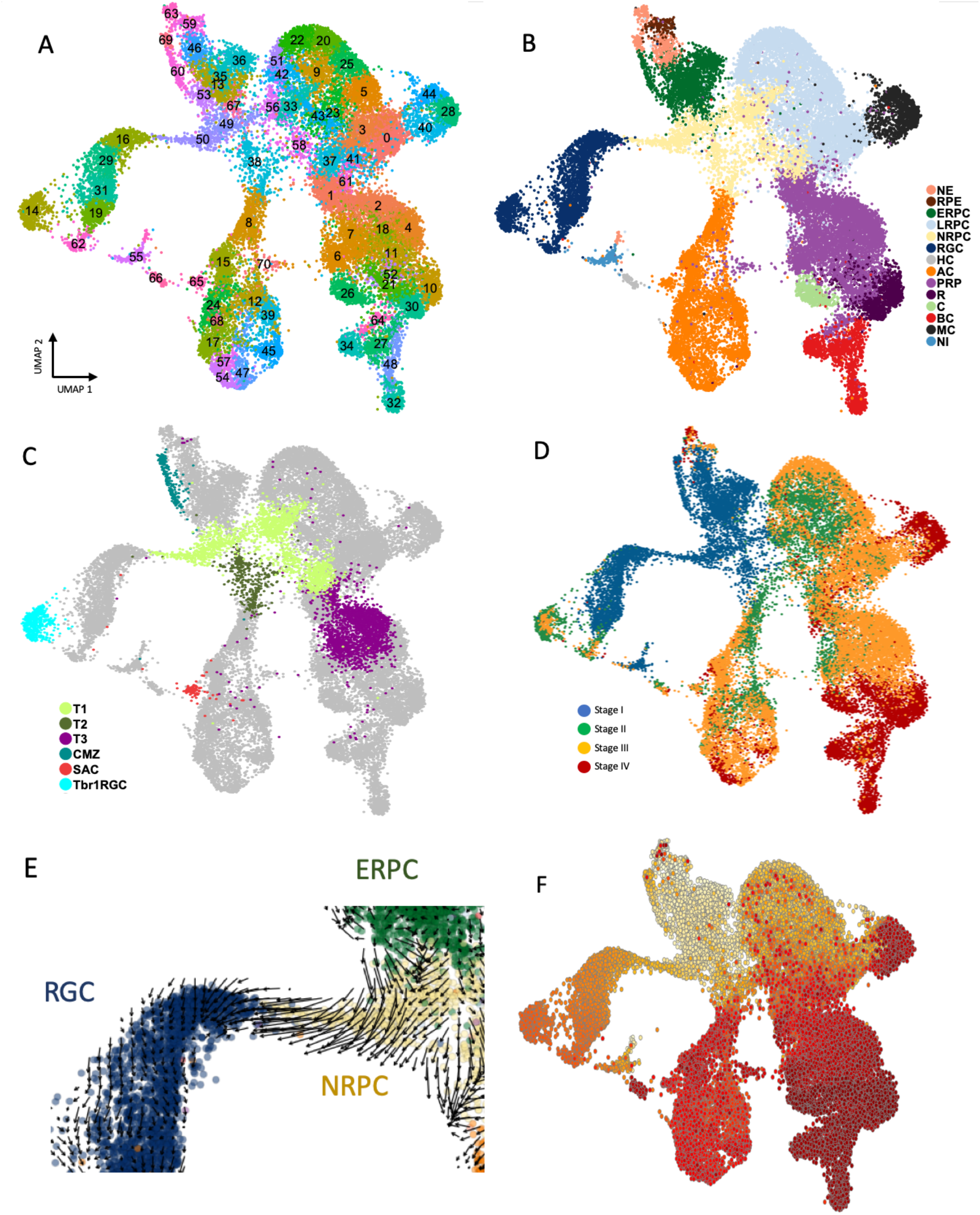
Joint scRNA-Seq-based UMAP of 38,091 cells corresponding to four developmental stages of native (NaR) and iPS-derived (3D-RA) murine retina. **(A)** 2D UMAP manifold showing NaR and 3D-RA cells jointly and their assignment to 71 clusters by k-means clustering. **(B)** Merging of the clusters in 13 major retinal cell types corresponding to neuroepithelium (NE), retinal pigmented epithelium (RPE), early (ERPC), late (LRPC), neurogenic retinal progenitor cells (NRPC), retinal ganglionic cells (RGC), horizontal cells (HC), amacrine cells (AC), photoreceptor precursor cells (PRP), cones (C), rods (R), bipolar cells (BC), and Müller cells (MC), on the basis of the expression of known marker genes (SFig. 2). **(C)** Identifying known retinal sub-populations: post-mitotic transitional precursor cell populations (T1, T2, T3)[16], Ciliary Marginal Zone (CMZ)[27], Tbr1+ retinal ganglionic cells (Trb1RGC)[28], and starburst amacrine cells (SAC)[29]. **(D)** Cells colored by developmental stage: I. blue = DD13 + E13, II. green = DD21 + P0, III. orange = DD25 + P5, IV. red = DD29 + P9. **(E)** Cell-specific RNA velocities [11] confirming the ERPC -> NRPC (T1) -> RGC cellular trajectory. **(F)** Velocity pseudo-time analysis using a velocity-inferred transition matrix [12]. Increase in pseudo-time is marked by increase in redness.

We defined gene expression signatures for 13 recognized retinal cell types using published information [25] (STable 3 and SFig. 2), and regrouped the clusters accordingly in 13 cell types corresponding to neuroepithelium (NE), retinal pigmented epithelium (RPE), early (ERPC), late (LRPC), and neurogenic retinal progenitor cells (NRPC), retinal ganglion cells (RGC), horizontal cells (HC), amacrine cells (AC), photoreceptor precursor cells (PRP), cones (C), rods (R), bipolar cells (BC), and Müller cells (MC) (Fig. 1B). Using additional gene expression signatures we further identified: (i) actively dividing ERPC, LRPC and NRPC (in S and G2-M phases of the cell cycle)[26], (ii) T1, T2 and T3 post-mitotic transitional precursor cell populations recognized in human native and hPSC-derived retina [16], (iii) the ciliary marginal zone (CMZ) [27], (iv) a recently described subgroup of Tbr1^+^ RGC cells located in the inner plexiform layer [28], and (v) starburst AC [29](Fig. 1C, SFig. 2 and STable 3).

Labelling cells by developmental stage (stages I to IV) distinguished ERPC from LRPC, and revealed the expected sequence of emergence of RGC (stage I), followed by HC, AC and PRP (stage II and III), then C, R, BC and MC (stage III and IV). Cells assigned to the Tbr1+ RGC cluster appeared at stage II and III. T1, T2 and T3 cells appeared in that order, and starburst AC at stage II and III (Fig. 1D). The UMAP manifold connected cell types consistently with known developmental trajectories [16,25,30,31], including: (i) NE -> RPE, (ii) NE -> ERPC, (iii) ERPC -> NRPC (T1) -> RGC, (iv) LRPC -> NRPC (T1->T2) -> AC, (v) LRPC -> NRPC (T1) -> PRP (T3) -> C/R, and (vi) LRPC -> MC. Reminiscent of previous studies [16,25], the cluster of HC cells was disconnected from the rest of the manifold providing no information about their precursors. In agreement with [25], BC appeared to emerge from PRP cells distinct from NRPC or T3 (Suppl. Video: http://www.sig.hec.ulg.ac.be/giga). Cell-specific RNA velocities [11] were consistent with the ERPC -> NRPC -> RGC trajectory but otherwise difficult to interpret (Fig. 1E). However, velocity pseudotime analysis (using a velocity-inferred transition matrix) implemented with scvelo [12] was remarkably proficient at ordering the four stages of development, as well as at identifying terminal cellular states (without benefitting from any information about development stage or root cells)(Fig. 1F).

### Comparison of NaR and 3D-RA cell fates in UMAP space highlights commonalities and differences in developmental trajectories

We then focused on the comparison between the behavior of NaR and 3D-RA cells. Global comparison of the distribution of NaR and 3D-RA cells across the manifold indicates that in vitro neuro-retinal differentiation from iPSCs largely recapitulates native development (Fig. 2A). This is substantiated by noting that 82% of the 71 clusters and 86% of the 13 cell types contain at least 10% of the least represented cell origin (NaR or 3D-RA) (Fig. 2B&C).

**Figure 2:**
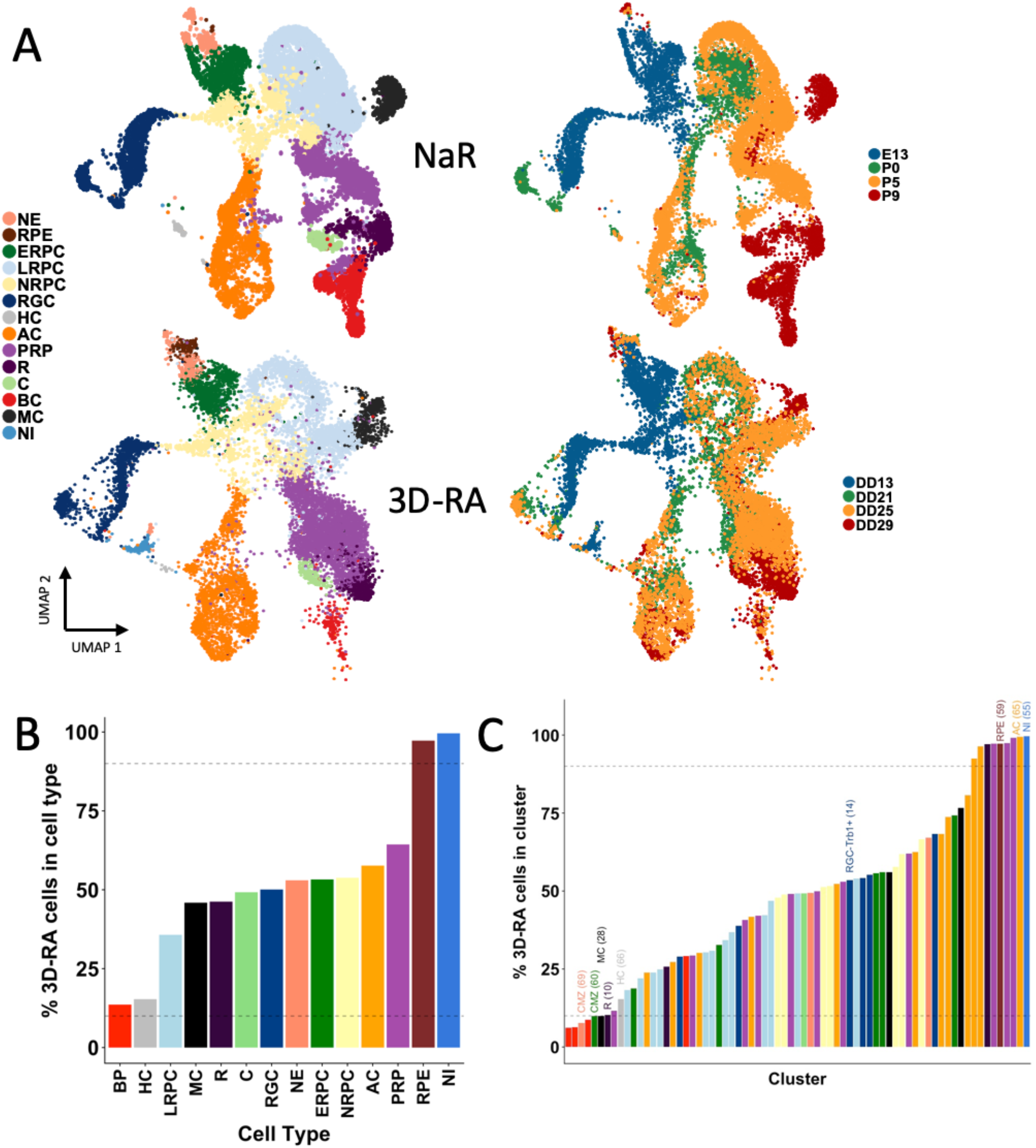
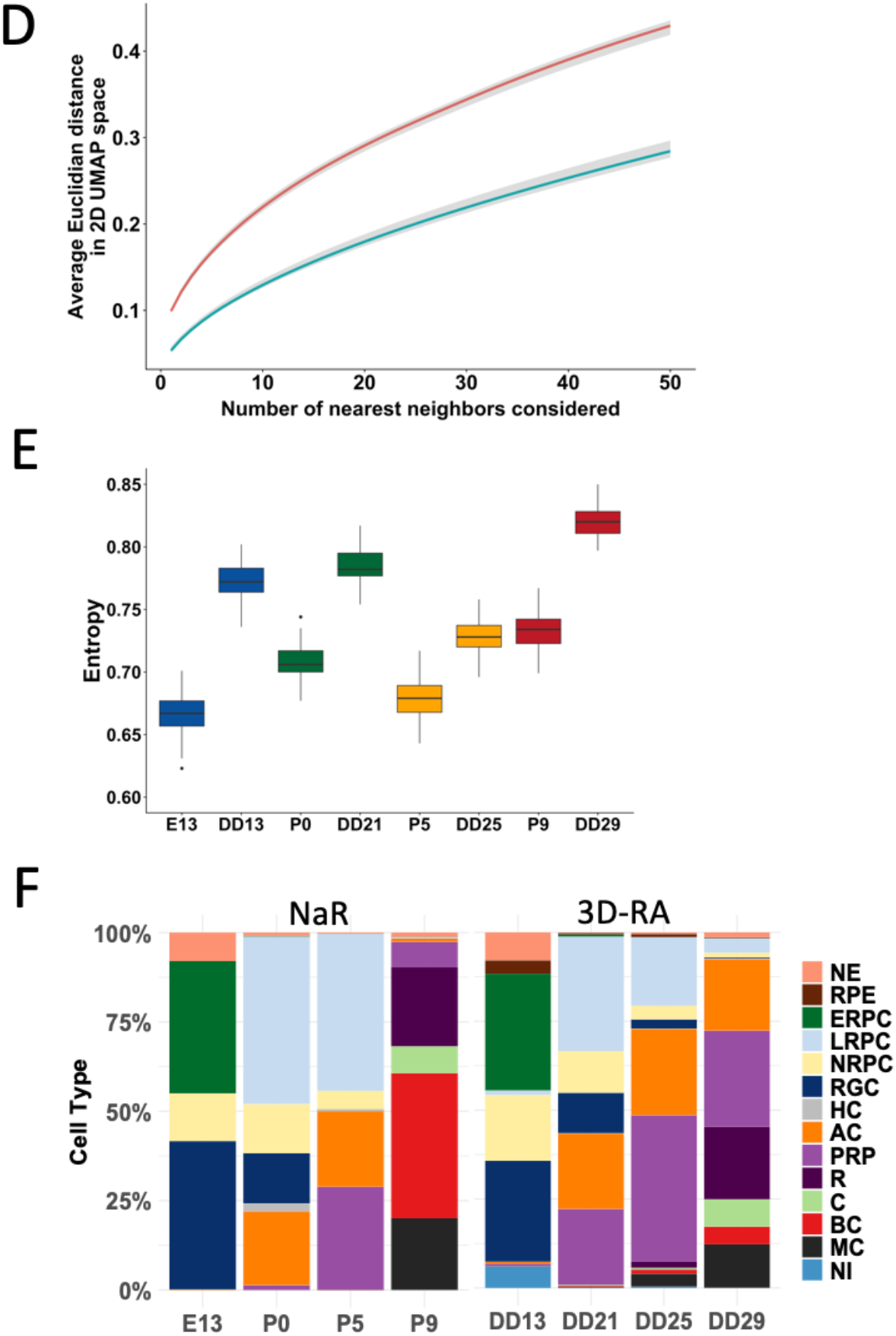
Comparison of Nar and 3D-RA cells in scRNA-Seq-based UMAP space. **(A)** Distribution of NaR (upper) versus 3D-RA (lower) cells across the UMAP manifold, sorted by cell type (left) and developmental stage (right). **(B-C)** Proportion of 3D-RA cells (adjusted for number of NaR and 3D-RA cells) in 14 cell types (B) and 71 clusters (C). 86% of cell types and 82% of clusters contain at least 10% of the least represented cell origin (NaR vs 3D-RA). Cell types are colored as in (A) and clusters are colored according to the cell type to which they were assigned. Notable clusters discussed in the main text are highlighted. Cluster 65 corresponds presumably to starburst AC. **(D)** Larger average distance in 2D UMAP space (Y-axis) from *n* nearest neighbors (X-axis) for 3D-RA (red) than for NaR cells (blue). **(E)** Larger cell type diversity (sampling-based measure of entropy) in the four developmental stages for 3D-RA than for NaR. **(F)** Proportions of cell types within developmental stage for NaR (left) and 3D-RA (right).

More granular examination, however, reveals noteworthy differences. The first one is the occurrence of NaR- or 3D-RA specific clusters and cell types: (i) the RPE cell type is almost exclusively composed of 3D-RA cells as a result of RPE elimination from NaR by dissection; (ii) the CMZ is absent in 3D-RA (only recently were culture conditions established for inducing selective CM retinal differentiation in human iPSC-derived RA [32]); (iii) AC cluster 65, thought to correspond to starburst AC, was only observed in 3D-RA; (iv) BC clusters 32, 34 and 48 are nearly exclusively composed of NaR cells, and (v) cluster 55 is exclusively populated by 3D-RA cells. Cluster 55 is thought to result from aberrant in vitro differentiation of NE into non-retinal neuronal cells. Indeed, it is connected to NE by a cellular bridge (Video: http://www.sig.hec.ulg.ac.be/giga), and strongly expresses Tbr1 and other genes typical of developing cortical neurons including reelin (STable 4&6). It is therefore

The second difference is the apparent relaxation of pseudo-spatial and pseudo-temporal transcriptome control in 3D-RA versus NaR. The developmental pathways traversed by NaR cells indeed appear tighter than those of 3D-RA cells, while NaR cells sampled at a specific developmental stage seem to populate fewer cell types than 3D-RA cells. To quantify the former, we down-sampled cells to equalize NaR and 3D-RA numbers (within developmental stage) and computed the average distance from the *n* closest neighbors, which was indeed highly significantly shorter for NaR than for 3D-RA (Fig. 2D). To quantify the latter, we measured the diversity of cell types within stages (using a measure of entropy), which was indeed significantly lower in NaR than in 3D-RA for all four stages (Fig. 2E). The last noteworthy differences between both systems is the observation that PRP arise earlier in 3D-RA than in NaR and accumulate at the expense of other cell types (particularly LRPC), yet partially fail terminal differentiation particularly into BC cells (Fig. 2A&F).

### 3D-RA culture conditions perturb genes and pathways that play key roles in NaR development

To identify key genes for retinal differentiation, we performed differential expression analysis for each cell type against all others, first considering NaR cells only. In NaR, we identified a total of 4,177 genes with significantly higher expression in at least one of the 13 main cell-types (as defined above) compared to all other cell types merged (log-fold change ≥ 0.25 and p-value ≤ 0.001), hereafter referred to as “cell type-specifying” genes (Fig. 3A and STable 4). Of those, 3,675 were also identified as dynamically regulated genes when using Monocle 2 [14] (SFig. 3 and STable 5).

**Figure 3:**
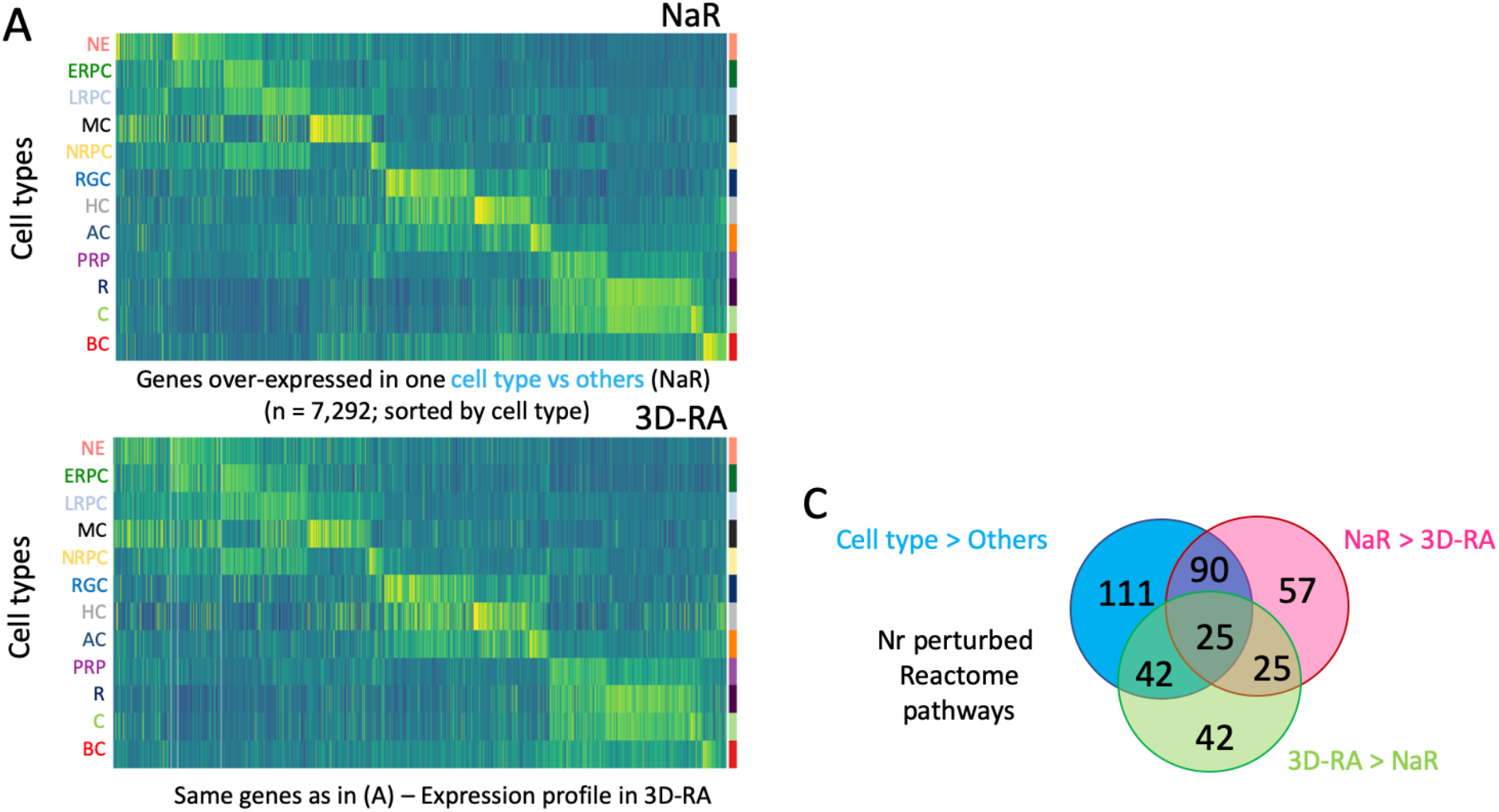
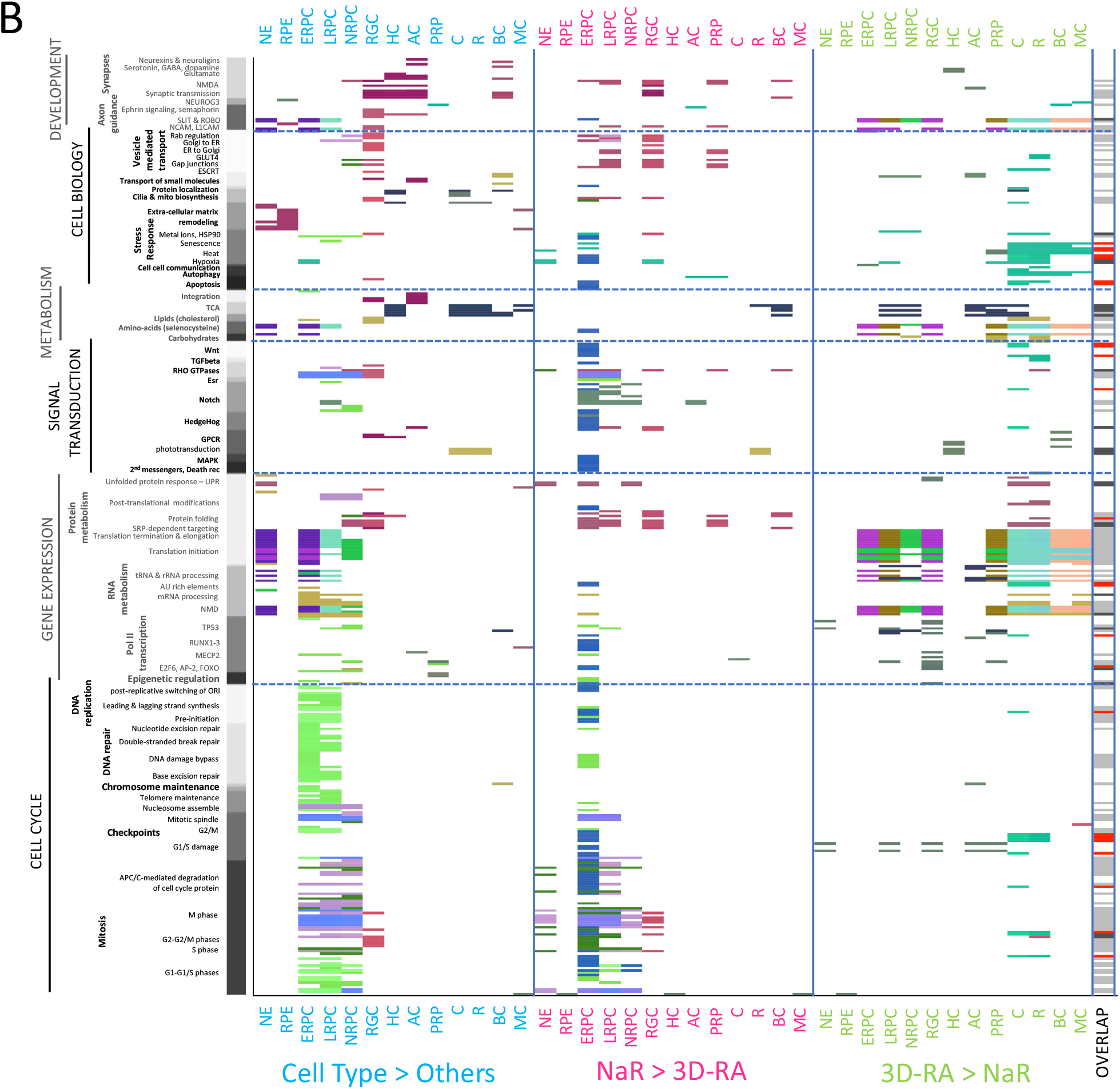
Comparison of the cell type-specific transcriptome of NaR and 3D-RA by means of scRNA-Seq. **(A)** Expression profiles in 12 cell types of 7,292 genes that are dynamically regulated during in vivo retinal development (i.e. significantly overexpressed in at least one cell type when compared to all other ones in NaR) in NaR (upper panel) and 3D-RA (lower panel). Abbreviations refer to cell types and are as defined in Fig. 1 (including color code). **(B)** Reactome pathways that are significantly (p ≤ 0.001) enriched amongst differentially expressed genes (“Cell type > Other”: when comparing expression levels between specific cell types and all other cells in NaR only; “NaR > 3D-RA” and “3D-RA > NaR”: when comparing expression levels between NaR and 3D-RA cells within cell type). Y-axis: Reactome pathways sorted by “top level” system (cell cycle, gene expression, signal transduction, metabolism, cell biology and development) and sub-level therein. Tiles mark the pathways that are significantly enriched in the corresponding contrast and cell type. The colors of the tiles reflect similarity in “found entities” as described in the main text and SFig. 4. Last column (“Overlap”): White: pathways significant in one contrast only, Black: pathways significant in all three contrasts, Grey: pathways significant in “Cell type > Other” and (“NaR > 3D-RA” or “3D-RA > NaR”), Red: pathways significant in “NaR > 3D-RA” and “3D-RA > NaR”. **(C)** Number of unique and shared Reactome pathways between “Cell type > Other”, “NaR > 3D-RA” and “3D-RA > NaR”. All overlaps are highly significant (p < 10^−6^) assuming random sampling from 2,365 Reactome pathways.

We then searched for enriched Reactome pathways [33,34] in the 13 lists of “cell type-specifying” genes. Two hundred sixty-eight pathways were significantly enriched (q ≤ 0.01) in at least one cell-type (STable 6). These corresponded primarily to: (i) accelerated cell division in ERPC, LRPC and NRPCs, (ii) intense post-transcriptional and translational activity in NE, ERPC, LRPC and NRPCs, (iii) activation of RHO GTPase- and NOTCH-dependent signaling in ERPC, LRPC, NRPCs, RGC and LRPC, NRPCs, respectively, as well as the GPCR-dependent phototransduction cascade in C and R, (iv) activation of mitochondrial citric acid (TCA) cycle and respiratory electro transport in HC, C, R, BC, and MC, of cholesterol synthesis in ERPC and RGC, and of insulin- and glucagon-dependent metabolic integration in RGC and AC, (v) enhanced remodeling of the extracellular matrix in NE, RPE and MC, and GAP junction trafficking in RGC, and (vi) activation of ROBO receptors-dependent axon guidance in NE, ERPC and LRPC, and of synapse formation in RGC, HC, AC and BC (Fig. 3B).

A Reactome pathway is considered enriched (in a list of submitted genes) if the number of genes in the list that are part of the pathway (the number of “found-entities”) is higher than expected by chance alone [33,34]. The found-entities for different enriched Reactome pathways often show considerable overlap. As an example, the same six genes (*Rfc5;Rfc4;Rfc1;Rfc2;Prim1*) in the list of 465 ERPC-specifying genes explain the enrichment of the “Leading strand synthesis” and “Polymerase switching” Reactome pathways (STable 6). We devised a method to assign colors to sets of found-entities such that strongly overlapping sets would have similar colors, while non-overlapping sets would have distinct colors (Methods and SFig. 4). As an example, we can see from Fig. 3B that the 48 Reactome pathways highlighted in NE correspond to six distinct sets of found entities (six dominant colors), that one of these sets is also driving Reactome pathway enrichment in RPE (bordeau), and that two others (indigo blue and purple) are also driving pathway enrichment in ERPC.

At first sight, genes that were differentially expressed between cell-types in NaR appeared to recapitulate their in vivo expression profile quite well in 3D-RA (Fig. 3A). Yet, to better appreciate the differences between in vivo and in vitro retinal differentiation, we performed differential expression analysis between NaR and 3D-RA separately for each cell type. For each of the 13 major cell types, we generated two lists of genes corresponding respectively to genes that were under-expressed in 3D-RA when compared to NaR (NaR>3D-RA) and genes that were over-expressed in 3D-RA when compared to NaR (3D-RA>NaR) (q ≤ 0.01; STable 7). We then searched for biological pathways that were over-represented in the corresponding gene lists using Reactome. This yielded 197 downregulated (NaR > 3D-RA) and 134 upregulated (3D-RA > NaR) pathways (Fig. 3B and STable 8). Strikingly, both down- and upregulated pathways (i.e. when comparing NaR and 3D-RA by cell type) exhibited considerable overlap with the pathways identified (see previous paragraph) when comparing cell types within NaR (Cell type > Others) (115/197, p < 10^−6^ and 67/134, p < 10^−6^) (Fig. 3C). More specifically, (i) the rate of cell division in NE, ERPC, LRPC and NRPC was reduced in 3D-RA when compared to NaR, (ii) post-transcriptional and translational mechanisms were exacerbated in ERPC, LRPC, NRPC, RGC, PRP, C, R, BC and MC of 3D-RA, when compared to NaR, (iii) signal transduction via WNT, TGF-beta, RHO GTPases, Esr, Notch, Hedgehog, MAPK, and Death receptors was diminished in 3D-RA when compared to NaR, particularly in ERPC and LRPC, while the phototransduction cascade was less active in 3D-RA-derived R than in NaR-derived R, (iv) mitochondrial citric acid (TCA) cycle and respiratory electron transport was increased in 3D-RA’s LRPC, NRPC, AC, PRP and C (yet increased in BC), cholesterol synthesis increased in 3D-RA’s C and R, and gluconeogenesis increased in 3D-RA’s PCP and R, (v) stress response and apoptosis was reduced in 3D-RA’s ERPC, yet increased in 3D-RA’s C, R, BC and MC (i.e. at the latest stages of 3D-RA culture), and (vi) vesicle mediated transport and synapse formation was decreased in 3D-RA’s LRPC, RGC and PRP (Fig. 3B). As testified by their assigned colors in Fig. 3B, the found-entities driving Reactome pathway enrichment when analyzing cell-type specifying genes (Cell type > Others) or when comparing NaR and 3D-RA (NaR > 3D-RA and 3D-RA > NaR) showed considerable overlap (see also SFig. 4). Thus, the genes and pathways that appear to be the most perturbed by the 3D-RA culture conditions are also the ones that play key roles in NaR development (i.e. the cell type-specifying genes as defined above).

### The expression level of many transcription factors is perturbed in 3D-RA

The 4,177 cell type-specifying genes in NaR (i.e. Cell type > Others, cfr above) comprised 293 transcription factors (TF)[35], including 107 that were at least 1.5 times more strongly expressed in one cell type when compared to any of the other cell types (Fig. 4A and STable 4). The latter comprised 88 TF that were previously reported in the context of retinal development, as well as 19 novel ones (NE: Peg3; LRPC: Lrrfip1; MC: Creb3l2, Csrnp1, Dbp, Nr4a1, Nr4a3; HC: Zfp618, Zfp804a; AC: Zfp503; PRP: Foxo3, Lcorl; R: Zfp516, Trps1, Ppard, Zc3h3, Mier1, Mier2, Lyar; BC: St18) (STable 9). Contrary to the overall expression profile (Fig. 3A), visual examination of the expression profiles of the 104 most differentially expressed TF indicated considerable loss of cell-type specificity in 3D-RA (Fig. 4A). Indeed, 155 of the 293 (53%) differentially expressed TF were significantly (q < 0.01) under-expressed in at least one cell type in 3D-RA when compared to NaR, while 80/293 (27%) were significantly (q < 0.01) over-expressed in at least one cell type (Fig. 4B and SFig. 6). Striking examples include Skil (ERPC), HevL (LRPC), Neurog2 (NRPC), Lhx1 (HC), Neurod2 (AC), Insm2 (PRP), Nfic (C/R), Ahr (C/R), Bhlhe23 (BC) and Nr1d1 (MC), which are all significantly under-expressed in 3D-RA when compared to NaR (Fig. 4C).

**Figure 4:**
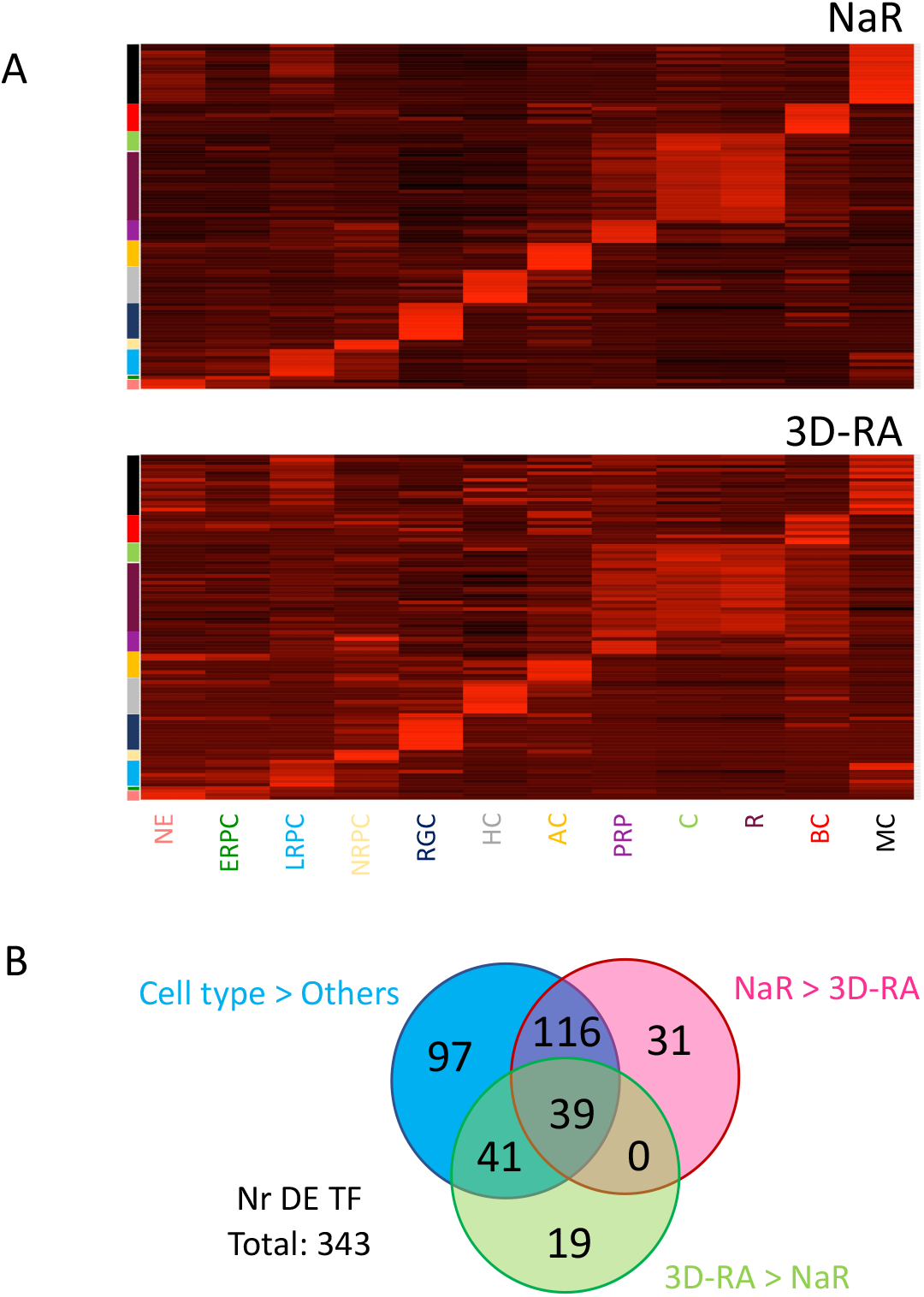
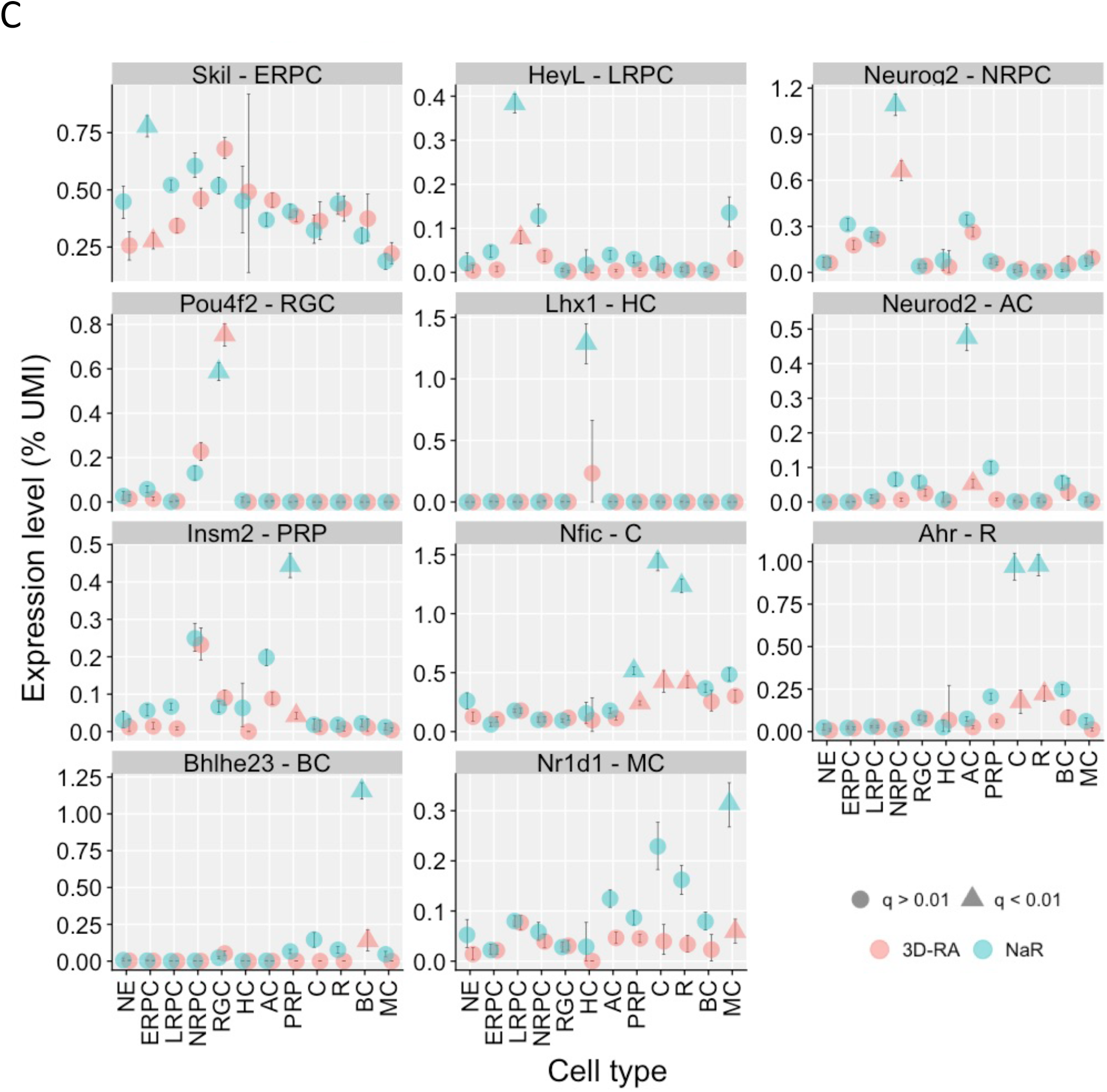
Comparison of the cell type-specific expression levels of TF in NaR and 3D-RA by means of scRNA-Seq. **(A)** Standardized expression levels of 104 most cell type-specific TF across 12 cell types in NaR (upper panel) and 3D-RA (lower panel). Abbreviations refer to cell types and are as defined in Fig. 1 (including color code). **(B)** Number of differentially expressed TF in “Cell type > Others”, “NaR > 3D-RA”, and “3D-RA > NaR”, with corresponding overlaps. The overlaps are highly significant (p < 10^−6^) assuming that TF are sampled randomly from the full collection of ∼1,500 TFs [35]. **(C)** Examples of TF that are (i) significantly overexpressed in one cell type when compared to all others in NaR, and (ii) significantly under- or over-expressed in that cell type between NaR and 3D-RA. The average expression levels (fraction of UMI) of the corresponding genes in the different cell types are shown for NaR (green) and 3D-RA (red). The error bars correspond to 99% confidence intervals determined by bootstrapping (n=1000). Green triangles mark cell types in which the corresponding gene is significantly (q < 0.01, i.e. accounting for multiple testing) overexpressed in NaR when compared to all other cell types combined. Red triangles mark cell types in which the expression level differs significantly (q < 0.01) between NaR and 3D-RA. The gene name and cell type of interest are given in the facet headers.

An additional 31 TF (not part of the list of cell type-specifying genes) were down-regulated in 3D-RA, while 19 were upregulated (SFig.6). Thus, the expression profile of a remarkably high number of TF appears perturbed in 3D-RA, and this may in part drive the differences observed between both systems, including with regards to Reactome pathways.

### Combined analysis of scRNA-Seq and bulk ATAC-Seq data reveals putative regulatory toggles in NaR

It is generally assumed that execution of the transcriptional program underlying differentiation is controlled by dynamically regulated TF that activate downstream target genes. To verify this assertion, we first performed bulk ATAC-Seq [36] on the first three stages of NaR (E13, P0, P5) and 3D-RA (DD13, DD21, DD25) samples to identify gene-switch components accessible during retinal development based on chromatin openness (SFig. 7A). For each sample type, we analyzed two technical replicates of two biological replicates for a total of 24 libraries. We defined a total of 123,482 peaks using MACS2 [37] (STable 10). Of these, 93,386 (75.6%) were detected in NaR, 97,333 (78.8%) in 3D-RA. 18,933 (15.3%) were common to all samples, 26,149 (30.0%) NaR-specific, 30,096 (24.4%) 3D-RA-specific, and 4,703 developmental stage-specific (3.8%; stage I: 294, stage II: 82, stage III: 4,327). The number of peaks increased with developmental stage in NaR but not in 3D-RA (highest number of peaks in DD13) (SFig. 7B). Nevertheless, stage I samples (E13 and DD13) clustered together, while for subsequent stages samples clustered by origin (NaR vs 3D-RA) (SFig. 7C). DNA binding motifs are reported for 151 of the TF found to be cell type-specifying by scRNA-Seq (see above), amounting to a total of 336 motifs (average number of motifs per TF: 2.3; range: 1 - 14). We used Homer to annotate our catalogue of ATAC-Seq peaks for the corresponding motifs [38]. In total Homer identified 7,128,225 binding motifs in 98,181 ATAC-seak peaks assigned (based on closest proximity) to 19,171 genes (STable 11).

To test whether TF that were overexpressed in a given cell type were indeed activating downstream target genes (as expected for “activator” TF), we searched for an enrichment of the cognate binding motifs in the ATAC-Seq peaks of genes that were significantly over-expressed in that cell type (relative to ATAC-Seq peaks of genes that were significantly under-expressed in the same cell type). As an example, the Crx TF is overexpressed in PRP, C and R: are ATAC-Seq peaks in the vicinity of the genes that are overexpressed in these cell types enriched in Crx binding motifs as expected if Crx is an activator TF? We used average number of binding motifs per ATAC-Seq peak per gene (total number of binding motifs divided by number of peaks) as metric to correct for gene length. We first analyzed NaR, and found 84 instances of binding motif enrichment (q-value < 0.01) for 37 TF over-expressed in the corresponding cell types (Fig. 5A, STable 12&13). Examples of such activator TF include Crx (enrichment of all three known Crx binding motifs: Crx-1, Crx-2 and Crx-3) in PRP, R and C, and Etv5 (1/1 binding motif) in LRPC (Fig. 5B).

**Figure 5:**
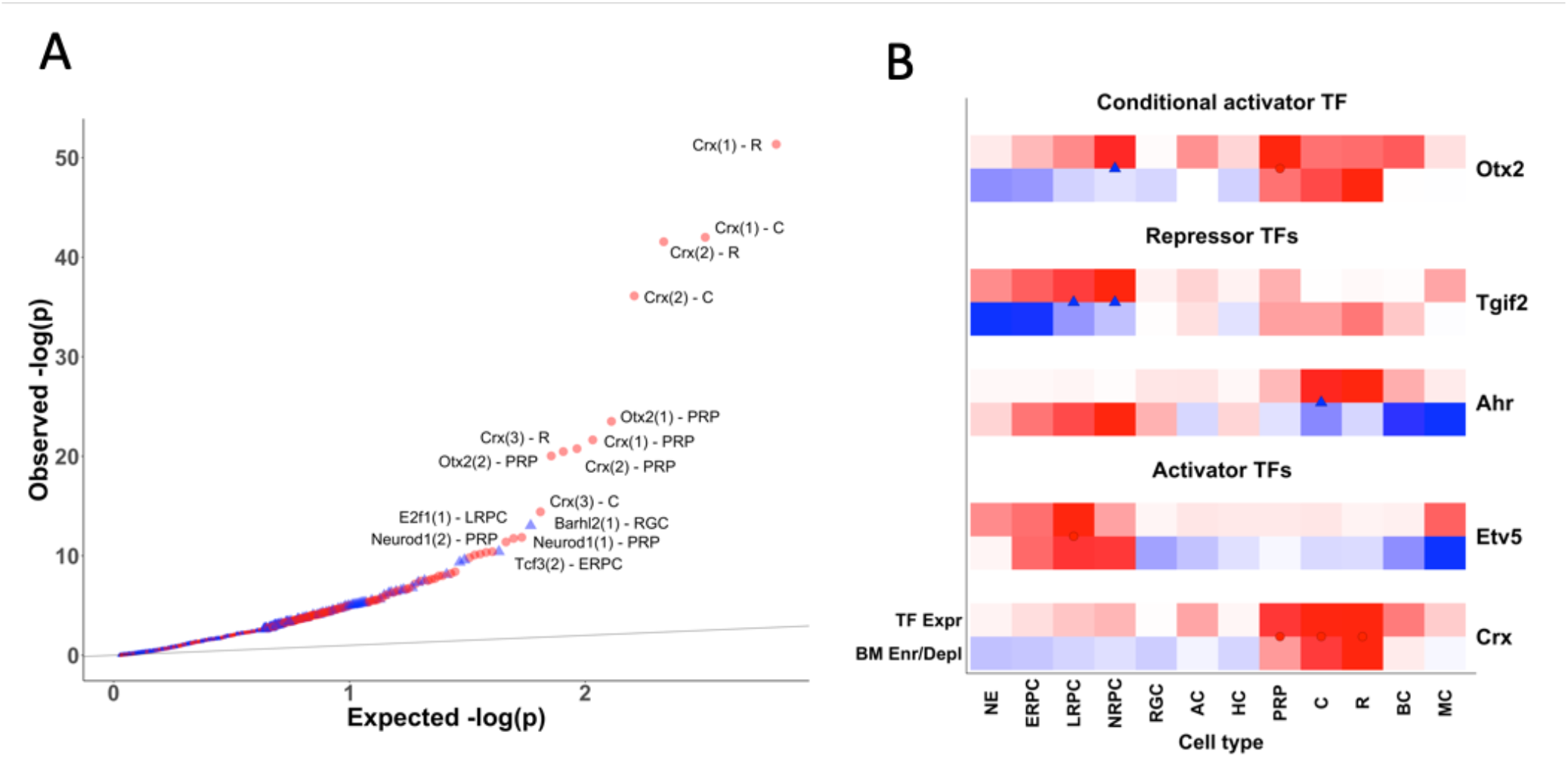
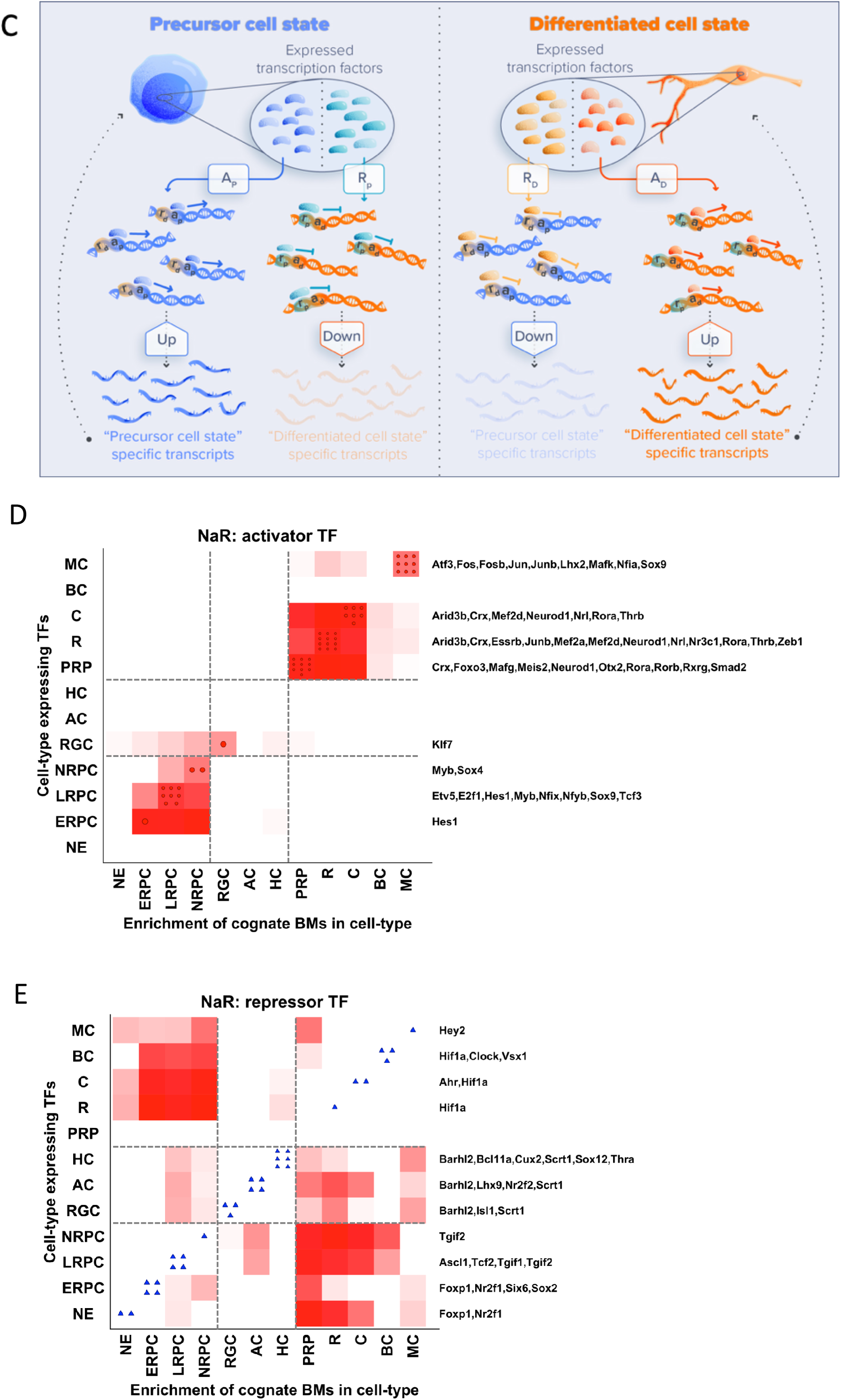
Combined analysis of scRNA-Seq and bulk ATAC-Seq data reveals putative activator and repressor TF that may constitute regulatory toggles. **(A) QQ plot of log(1/p) values of the difference in average number of binding motifs in ATAC-Seq peaks of over-expressed versus under-expressed genes for TF over-expressed in the corresponding cell type**. Enrichments are indicated by red dots, depletions by blue triangles. Symbols are large for TF with q-value < 0.01, and small for TF with q-value ≥ 0.01. For the most significant effects, the name of the TF is given, as well as – between brackets - the index of the binding motif, and the cell type. The grey line corresponds to expectations assuming that all tests are true null hypotheses. **(B) Examples of activator, repressor and conditional activator TF**. Upper lines (“TF Expr”, white-red color code): standardized expression pattern of corresponding TF across 12 cell types. Lower lines (“BM Enr/Depl”, blue-white-red color code): standardized enrichment (red) or depletion (blue) of binding motif(s) of corresponding TF across 12 cell types. The red circles and blue triangles mark the cases that are part of (A). They require that the TF be overexpressed in the corresponding cell type, and that there is either a significant enrichment of binding motifs in overexpressed genes (red circles) or a significant depletion (blue triangles). **(C) Components of regulatory toggles and principles underlying their detection**. Shown are a hypothetical precursor (blue) and derived differentiated cell (orange). The precursor cell is expressing a number of activator (A_P_, blue) and repressor TF (R_P_, green). These are respectively activating and repressing target genes by binding to motifs in cis-acting regulatory elements labelled respectively as a_P_ and r_P_. The differentiated cell is expressing its own activator (A_D_, orange) and repressor TF (R_D_, yellow), which are respectively activating and repressing target genes by binding to motifs in cis-acting regulatory elements labelled respectively as a_D_ and r_D_. Genes that are overexpressed in precursor cells (under-expressed in differentiated cells) are enriched in a_P_ and r_D_ binding motifs, and depleted in a_D_ and r_P_ binding motifs. Genes that are overexpressed in differentiated cells (under-expressed in precursor cells) are enriched in a_D_ and r_P_ binding motifs, and depleted in a_P_ and r_D_ binding motifs. **(D-E) Combined enrichment profile across all cell types (X-axis) of binding motifs for all activator (upper graph) and repressor (lower graph) TF expressed in a given cell type (Y-axis) in NaR**. Standardized (across entire array) sum of signed (+ for enrichment, - for depletion) log(1/p) values for binding motifs of TF expressed in a given cell type (Y-axis). Positive values are measured by a white-red color code; values ≤ 0 are in white. For each cell type, the number of overexpressed activator TF (upper graph, red circles on diagonal) and repressor TF (lower graph, blue triangles on diagonal) are given, and their names provided on the right. The horizontal and vertical dotted lines delineate clusters I, II and III as defined in the main text.

Intriguingly, we observed 45 instances of binding motif depletion (q-value < 0.01) for 26 other TF. Stated otherwise, for 26 TF that were over-expressed in a given cell type, binding motifs were significantly more abundant in ATAC-Seq peaks of genes that were under-expressed than in the ATAC-Seq peaks of genes that were over-expressed in the corresponding cell type (Fig. 5A, STable 12&13). One explanation of this finding is that these TF act as repressors rather than activators. Examples of such candidate “repressor” TF are Arh (1/2 binding motifs) in C, and Tgif2 (2/3 binding motifs) in LRPC and NRPC (Fig. 5B).

We reasoned that such repressor TF may be components of regulatory toggles that ensure cell type-specific gene expression not only by inducing expression of required genes (via activator TF), but also by precluding expression of undesired genes (via repressor TF) (Fig. 5C). To gain insights in the nature of the genes targeted by these putative repressor TF, we searched for cell types in which the corresponding binding motifs were enriched in over-expressed genes (even if the TF itself is not strongly expressed in that cell type). Fig. 5E shows the corresponding results for all repressor TF over-expressed in a given cell type jointly. Thus, for all candidate repressor TF identified in a given cell type (f.i. Foxp1, Nr2f1, Six6 and Sox2 in ERPC; see Fig. 5E), we computed the log(1/p) value of the difference in the density of binding motifs in over-versus under-expressed genes in each cell type, signed them according to the direction of the difference (enrichment versus depletion in over-expressed genes), and averaged these values across the (f.i. four in ERPC) candidate repressor TF. For comparison, similar plots (Fig. 5D) are shown for the combined effect of all activator TF expressed in a given cell type. This analysis revealed that the identified repressor TF systematically target genes that are overexpressed in (and hence specify) another retinal cell type than the one(s) in which they are expressed, with a clear pattern. It appears that the 12 cell types analyzed in NaR form three clusters: (I) NE, ERPC, LRPC and NRPC, (II) RGC, AC and HC, and (III) PRP, C, R, B, C and MC. Repressor TF which are expressed in cluster (I) are primarily targeting genes that are over-expressed in cluster (III), repressor TF which are expressed in cluster (II) are primarily targeting genes that are expressed in cluster (I) or (III), and repressor TF which are expressed in cluster (III) primarily target genes that are expressed in cluster (I). There was considerable overlap between the TF (activator and repressor) over-expressed in cell types from the same cluster. As an example, Hif1A is over-expressed in R, C and BC (cluster III), while Bahrl2 is over-expressed in RGC, AC and HC (cluster II)(Fig. 5E).

We also identified six “conditional activator” TF. These were characterized by the enrichment of binding motifs in over-expressed genes for some cell type(s), yet the depletion of binding motifs in over-expressed genes for other cell type(s) (Etv1, Fosb, Otx2, Pax6, Tcf3, Sox11) (Fig. 5B; STable 12&13). A good example is Otx2, which is significantly over-expressed in PRP and NRPC, and whose three binding motifs are enriched in genes that are over-expressed in PRP while being enriched in genes that are under-expressed in NRPC (which include the genes that are over-expressed in PRP). One possible explanation of these observations is that the corresponding TF are necessary but not sufficient to induce expression of target genes. As an example, Otx2 may be induced in NRPC that will develop into PRP, but only exert its transcriptional effects after maturation into PRP. Consistent with this hypothesis, NRPC expressing Otx2 tended to cluster in the vicinity of PRP in the UMAP manifold (SFig. 7). We therefore refer to these six TF as “conditional activators”.

Finally, three TF were characterized by the enrichment of one of their binding motifs (in overexpressed genes), yet the depletion of another of their binding motifs in the same cell type (Lhx1, Plagl1, Zic1) (STable 12&13). These will be referred to as “dual TF” yet were considered with caution.

### ScATAC-Seq supports the toggle hypothesis

To further test our toggle hypothesis, we took advantage of publicly available scATAC-Seq data for 1,792 cells isolated from retina of eight-week old mice (P56)[39]. Using 10xGenomics’ Cell Ranger ATAC software we clustered the cells based on transposase accessibility of 117,073 scATAC-Seq peaks. Clusters were then assigned to specific cell-types using the same gene signatures used with the scRNA-Seq data (STable 3). Cell-specific gene expression levels were estimated as the proportion of reads mapping to ATAC-Seq peaks assigned to the corresponding gene (i.e. the transposase accessibility of the gene in that cell). Consistent with the results reported in [39], these analyses confirmed that the sample is primarily composed of R (n = 736), AC (n = 500), BC (n = 419), C (n = 91), and MC (n = 46) (Fig. 6A).

**Figure 6:**
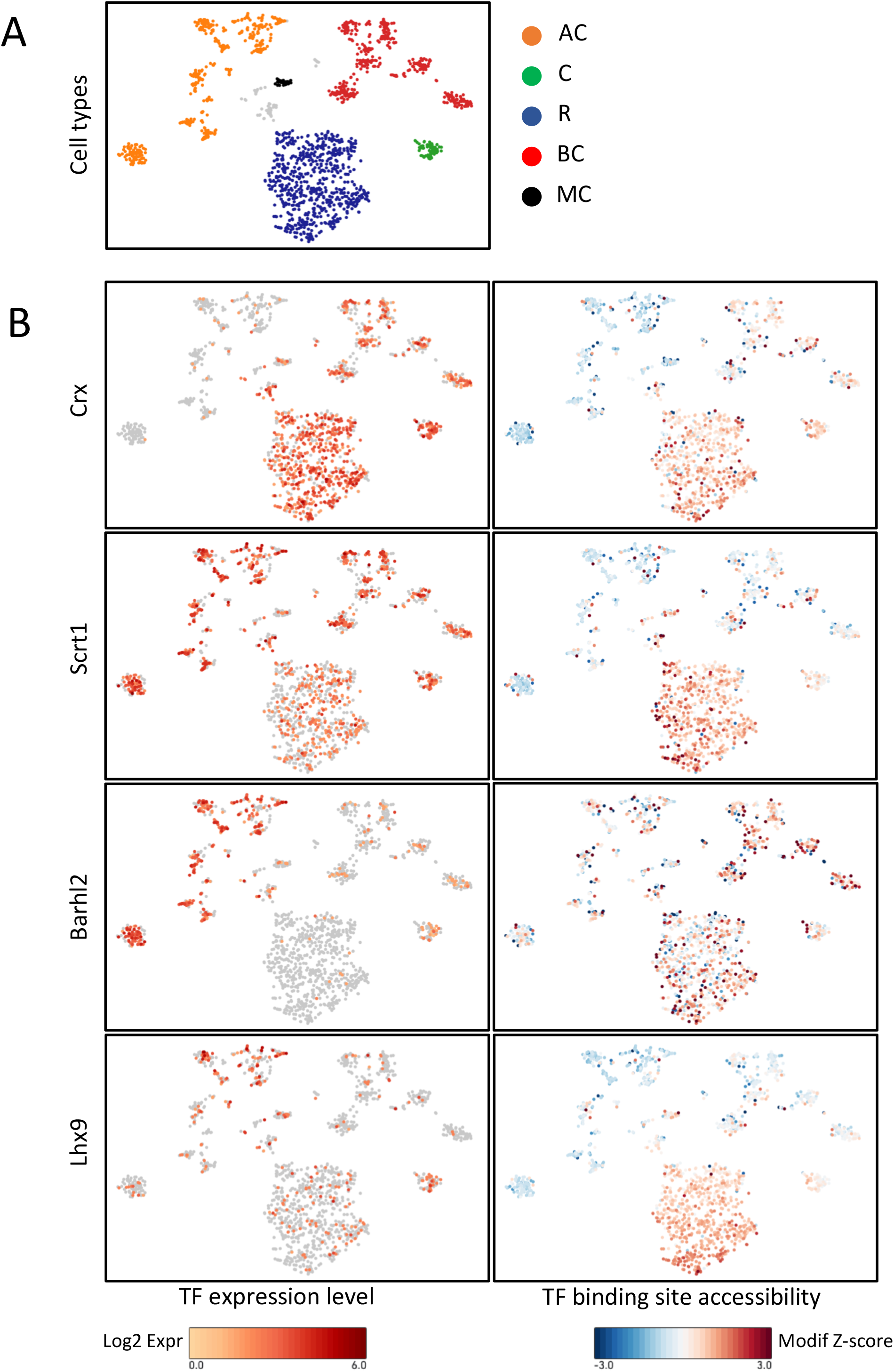
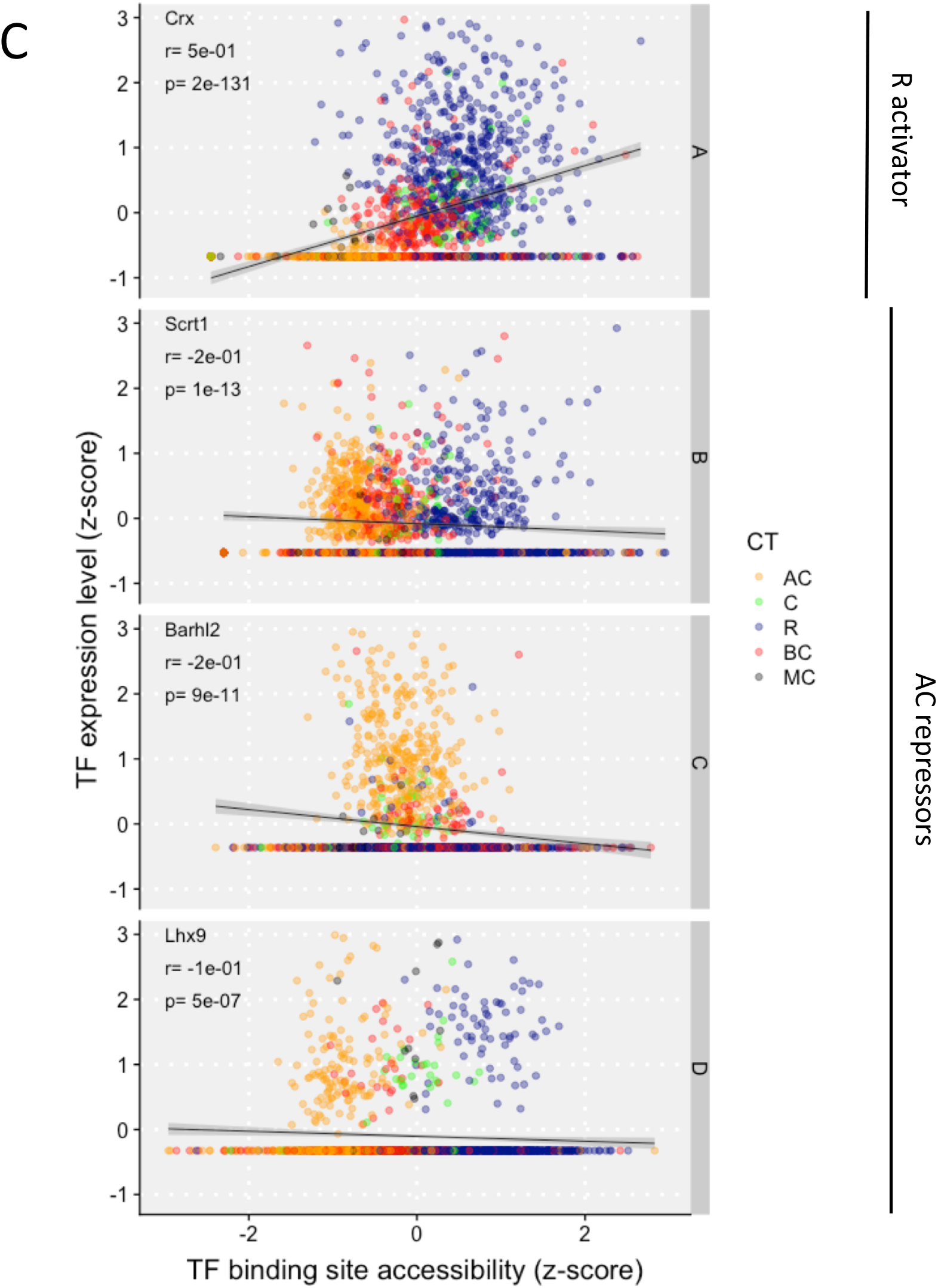
Single-cell ATAC-Seq data support the regulatory toggle model. **(A)** scATAC-Seq based tSNE plot constructed with the Cell Ranger ATAC software (10xGenomics) and the data from Norrie et al. [38]. Cells were assigned to cell-types based on the accessibility of the marker genes reported in STable 3. Cell-type abbreviations and colors are as in the previous figures. The number of cells per cell type are in good agreement with [39]. **(B)** (Left panels) Expression levels of one C/R activator (Crx) and three presumed AC repressors (Scrt1, Barhl2, Lhx9) inferred from their transposase accessibility, showing higher accessibility of Crx in R, C and BC, and higher accessibility of Scrt1, Barhl2 and Lhx9 in AC. (Right panels) Aggregate genome-wide transposase accessibility of cognate binding motifs, showing higher accessibility of Crx motifs in R, C and BC (i.e. same as for Crx itself), and higher accessibility of Scrt1, Barhl2 and Lhx9 motifs in R, C (and BC) than in AC (i.e. opposite as for corresponding TF). **(C)** Scatter plots showing the correlation between gene-specific TF transposase accessibility (y axis) and corresponding genome-wide TF binding motif transposase accessibility (x axis). Dots correspond to individual cells. All correlations are highly significant, positive for the C/R activator, and negative for the AC repressors. See SFig. 9 for plots of 11 C/R activators and 4 AC repressors.

Hence, this dataset provides scATAC-Seq information for one cell type belonging to cluster II (AC) and three cell types belonging to cluster III (R, C and BC). Examining the list of activator and repressor TF reported in Fig. 5D and E, we concluded that we could use this scATAC-Seq dataset to (i) test the predicted activator status of 11 of 12 TF active in C and/or R (i.e. expressed in C/R and activating genes in C/R) (Arid3b, Crx, Esrrb, Junb, Mef2a, Mef2d, Neurod1, Nrl, Nr3c1, Rora, Zeb1; no data available in Cell Ranger for Thrb; see Fig. 5D), and (ii) test the predicted repressor status of 4 out of 4 TF active in AC (i.e. expressed in AC and repressing genes in AC that are normally expressed in C/R)(Barhl2, Lhx9, Nr2f2, Scrt1; see Fig. 5E).

We first tested the 11 C/R activator TF. Cell-specific TF expression levels were estimated as the proportion of reads mapping to ATAC-Seq peaks assigned to the corresponding TF-encoding gene (i.e. the transposase accessibility of the TF-encoding gene in that cell). The aggregate expression levels of the genes activated by the TF were estimated as the proportion of reads mapping to genome-wide ATAC-Seq peaks matched to a cognate binding motif by the Cell Ranger ATAC algorithm (10X Genomics)(i.e. the aggregate accessibility of the binding motif of the TF of interest in that cell). Our prediction was that for C/R activator TF, both the TF-encoding genes and the cognate binding motifs (genome-wide) should be transposase-accessible in C and R cells (= cluster III). Visual examination of paired tSNE maps for (i) TF gene accessibility and (ii) cognate TF binding motif accessibility, clearly revealed the expected colocalization in C/R cells (as well as BC cells)(Fig. 6B). To more rigorously assess the statistical significance of these visual impressions, we computed the correlations between the cells’ TF gene accessibility and TF binding motif accessibility. The correlation was positive for 11/11 C/R activators (*p*_*Spearman*_ ≤ 0.05 for 9/11)(Fig. 6C and SFig. 9).

We then performed the same analyses for the four AC repressor TF. Visual examination of the paired tSNE plots revealed a striking contrast with the 11 activator TF: repressor TF gene accessibility was highest in AC (cluster II), while repressor TF binding motif accessibility was highest in the other cell types (cluster III) (Fig. 6B). Accordingly, the correlation between the cells’ TF gene accessibility and TF binding motif accessibility was negative for the four AC repressors (*p*_*Spearman*_ ≤ 0.05 for 4/4)(Fig. 6C and SFig. 9). The probability to observe a positive correlation for 11/11 predicted activator TF and a negative correlation for 4/4 predicted repressor by chance alone is 3×10^−5^.

### Comparing the effects of activator and repressor TF between NaR and 3D-RA using scRNA-Seq and bulk ATAC-Seq data

We repeated the same analyses as described above for the 13 major cell types detected in 3D-RA (including RPE). We identified 18 activator TF, 24 repressor TF, 3 conditional activator TF, and 2 dual TF (STable 12&14). Thirty-eight of these 47 TF overlapped with those found in NaR. Analyzing the combined effect of TF overexpressed in a given cell type largely confirmed the three clusters detected in NaR. RPE appear to be part of cluster (I), while 3D-RA MC could not be classified as no over-expressed TF (whether activator or repressor) could be detected in this cell type in 3D-RA (Fig. 8A&B). The degree of enrichment/depletion of TF binding motifs (measured by the signed (+ for enrichment, - for depletion) log(1/p)) in the different cell types was highly correlated between NaR and 3D-RA (r= 0.82, p < 2.2e-16)(Fig. 8C). The slope of the regression line was significantly < 1 suggesting that, overall, the TF’s activator and repressor effects might be slightly reduced in 3D-RA when compared to NaR. Alternatively, statistical power could be slightly higher in NaR due to the larger number of analyzed cells. Outliers included Crx and Otx2 which appeared to have more pronounced activator effects in PRP of 3D-RA than of NaR (Fig. 8C&D). This may be related to the fact that the 3D-RA culture conditions were designed to “push” PRP development (see also Fig. 2J). Other outliers were Etv1, Etv5, Hes1 and Zbtb7a which pointed towards TF-driven differences in the NaR and 3D-RA transcriptomes of HC and MC (Fig. 8C). Binding motifs for this group of TF were enriched in genes that were under-expressed in MC of NaR but this was not observed in 3D-RA (dots near horizontal dotted line Fig. 8C), while being enriched in genes that were under-expressed in HC of 3D-RA but this was not observed NaR (dots near vertical dotted line in Fig. 8C)(Fig. 8D). Of note, binding motifs of Etv1, Etv5, Zbtb7a, but not Hes1 are partially overlapping (SFig. 10). The prediction is therefore that (i) there are genes that are under-expressed in MC of NaR but not in MC of 3D-RA and whose ATAC-Seq peaks have a high density in Etv1/Etv5/Hes1/Zbtb7a binding motif, and (ii) there are genes that are under-expressed in HC of 3D-RA but not in HC of NaR and whose ATAC-Seq peaks have a high density in Etv1/Etv5,Hes1/Zbtb7a binding motif. Indeed, we found that there was a very strong coincidence between the density of Etv1/Etv5/Hes1/Zbtb7a binding motifs and the two predicted expression patterns (under-expressed in MC of NaR but not of 3D-RA; under-expressed in HC of 3D-RA but not of Nar)(Fig. 8E&F). We identified 77 genes under-expressed in MC of NaR but not of 3D-RA with high density in Etv1/Etv5/Hes1/Zbtb7a bindings motifs, and 61 genes under-expressed in HC of 3D-RA but not of Nar with high density in Etv1/Etv5/Hes1/Zbtb7a bindings motifs (STable 15). We submitted these gene lists to Reactome. The pathways that were enriched overlapped strongly between the two lists (although only eight genes were common to both lists) and pertained mainly to RNA stability and translation and to a lesser extent to the cell cycle (STable 15). They coincided remarkably well with the Reactome pathways that were highlighted by the genes that appeared in the scRNA-Seq analyses to be more strongly expressed in MC of 3D-RA than of NaR (found entities labelled in orange [3D-RA > NaR, column MC] in Fig. 3B). Thus, the combined scRNA-Seq and ATAC-Seq analysis reveals that part of the difference between MC of Nar and 3D-RA is due to perturbed expression of genes that are controlled by the Etv1/Etv5/Hes1/Zbtb7A TF squadron. Of note, these effects appeared largely independent of differences in the expression levels of the corresponding TF (in MC/HC) between NaR and 3D-RA per se: Etv1 was the only TF to have a significantly higher expression level in 3D-RA than NaR in MC (STable 7).

**Figure 8:**
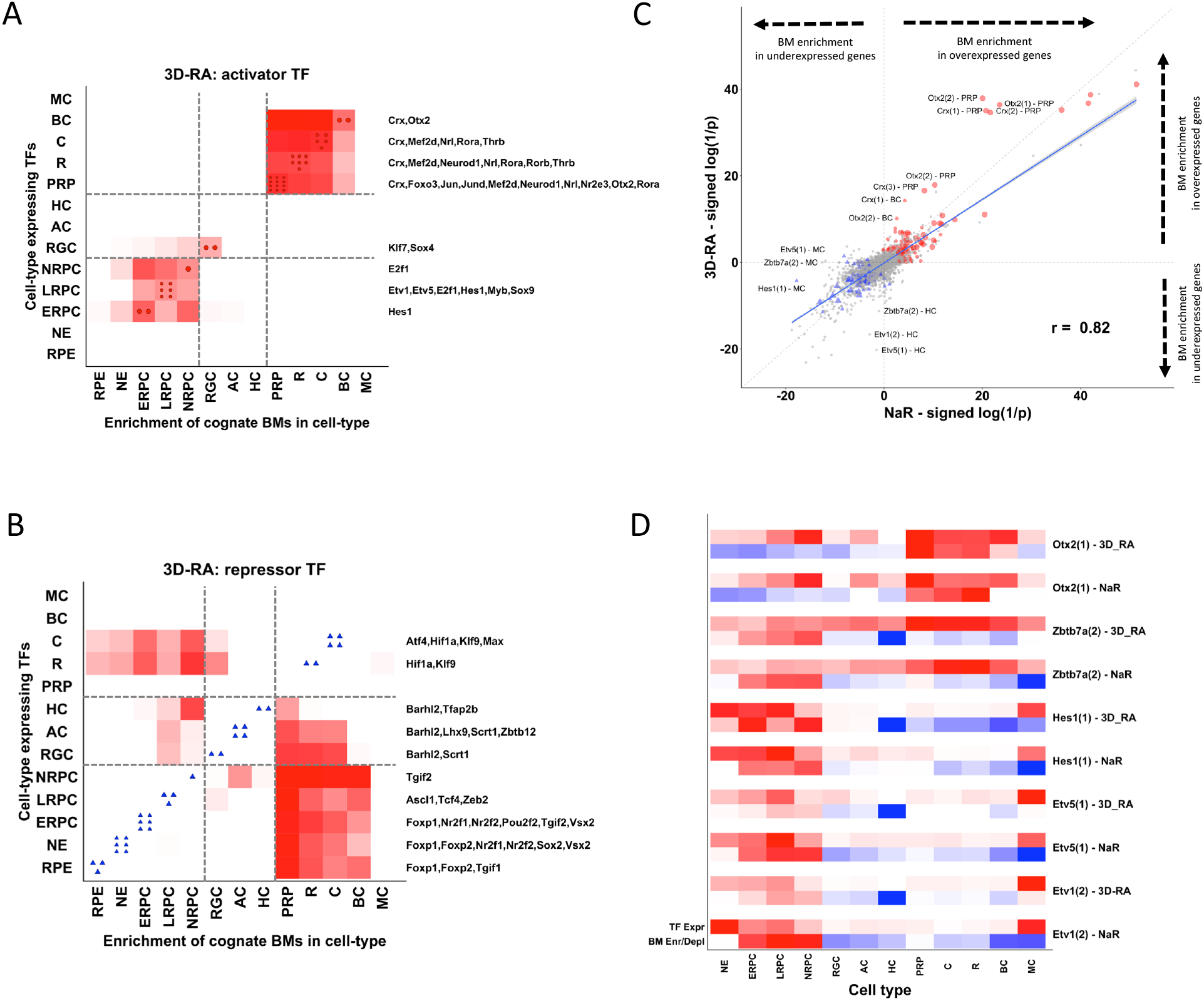
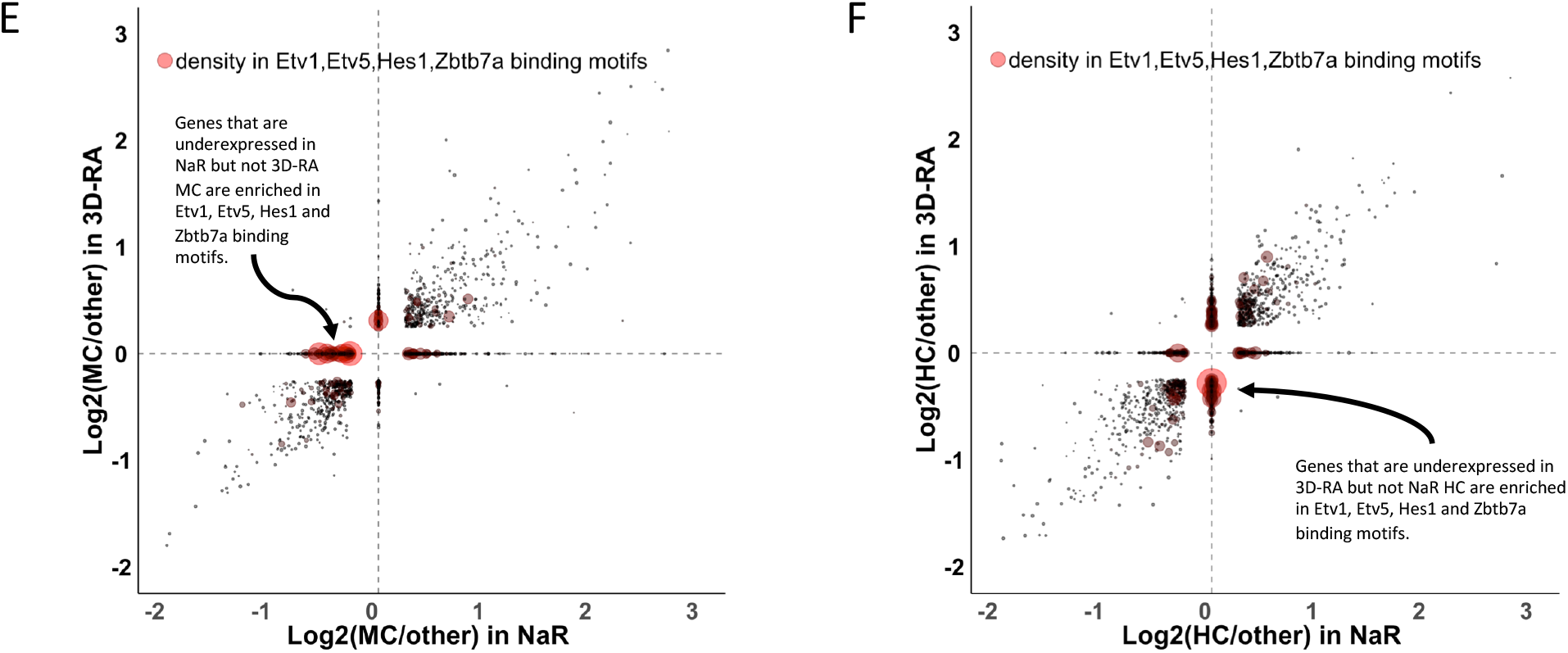
Comparing the operation of regulatory toggles between Nar and 3D-RA. **(A**,**B) Combined enrichment profile across all cell types (X-axis) of binding motifs for all activator (A) and repressor (B) TF expressed in a given cell type (Y-axis) in 3D-RA**. Standardized (across entire array) sum of signed (+ for enrichment, - for depletion) log(1/p) values for binding motifs of TF expressed in a given cell type (Y-axis). Positive values are measured by a white-red color code; values ≤ 0 are in white. For each cell type the number of overexpressed activator TF (upper graph, red circles on diagonal) and repressor TF (lower graph, blue triangles on diagonal) are given, and their names provided on the right. The horizontal and vertical dotted lines delineate clusters I, II and III as defined in the main text. **(C) Comparison of the log(1/p) values for enrichment (+)/depletion (-) of TF binding motifs across cell types between NaR (X-axis) and 3D-RA (Y-axis)**. Red circles: values for activator TF in the cell type in which they are overexpressed. Blue triangles: values for repressor TF in the cell type in which they are overexpressed. Small grey circles: values for cell types in which the corresponding TF is not overexpressed. The identity of outlier TF is given, as well as – between brackets - the index of the binding motif, and the cell type. “r” = correlation coefficient. **(D)** Expression levels (red scale) (upper line) and enrichment (red scale)/depletion (blue scale) of binding motifs in overexpressed genes (lower line) in NaR and 3D-RA for five TF highlighted in Fig. 8C. **(E)** Log2 fold over (+) or under (-) expression of genes in MC relative to the other cell types in NaR (x-axis) and 3D-RA (y-axis). When the fold-change in expression is not significantly different from 0, the gene receives 0 value in the corresponding sample type (i.e. on the horizontal line for 3D-RA, and on the vertical line for NaR). The size and redness of the symbols measures the density in the weighted sum of the binding motifs for Etv1, Etv5, Hes1 and Zbtb7a for the corresponding gene. One can clearly see that the genes that are significantly underexpressed in NaR but not in 3D-RA (genes on left side of horizontal line) are enriched in genes with high density of TF binding motifs. **(F)** As (E) for HC. One can clearly see that the genes that are significantly underexpressed in 3D-RA but not in NaR (genes on bottom side of vertical line) are enriched in genes with high density of TF binding motifs.

## Discussion

We herein use scRNA-seq to compare the unfolding of the epigenetic program in *in vivo* versus *in vitro* (from iPS cells) derived murine retina at four matched development stages encompassing the presumed emergence times of the major retinal cell types (E13 vs DD13, P0 vs DD21, P5 vs DD25 and P9 vs DD29). Results obtained by combining information from (i) the analysis of four developmental stages, (ii) 3D UMAP manifolds visualized in virtual reality (http://www.sig.hec.ulg.ac.be/giga), and (iii) RNA velocity analysis, are in good agreement with the previously reported, main retinal developmental trajectories (Fig. 1F). We identify >4,000 genes that are differentially expressed during *in vivo* retinal differentiation corresponding to tens of biological pathways pertaining to the cell cycle, gene expression, signal transduction, metabolism, cell biology and development (Fig. 3). Several of these pathways were previously highlighted when submitting differentially expressed genes identified from the analyses of bulk RNA-Seq data from multiple time points (E11 to P28) during retinal development [40]. Our data now allows to assign highlighted pathways to individual cell types. Differentially expressed genes include ∼300 TF, of which ∼100 are at least 1.5 times more strongly expressed in one specific retinal cell type when compared to all other ones. The latter include 19 TF not yet described in the field of retinal development which could serve as a starting point for functional investigations of the roles of these TF in retinogenesis and physiology.

We generated bulk ATAC-Seq data for three of the four analyzed developmental stages in both NaR and 3D-RA. This allowed us to identify 98,181 putative regulatory elements assigned (by proximity) to 19,171 genes, that are accessible during retinal development. This collection of ATAC-Seq peaks also allowed us to test the activity of 151 TF shown by scRNA-Seq to be differentially expressed during retinal development, and which have known binding motif(s). For 31 of these (considering both NaR and 3D-RA; STable 12), we observed an enrichment (q-value < 0.01) of binding motifs in ATAC-Seq peaks of genes that are over-expressed in the same cell type as the TF. This is what is expected for TF that act predominantly as activators in the corresponding cell type. Reassuringly, the list of predicted activators includes several TF that are known to play key roles during retinal development such as Crx, Neurod1, Nr2e3, Nrl, Rora, Rorb, Rxrg, Sox9 and Thrb [30, 41-43].

Unexpectedly, for 31 other TF we observed a significant depletion of binding motifs in the ATAC-Seq peaks of genes that were over-expressed in the same cell type as the TF. This is what is expected if the TF acts predominantly as a repressor in that cell type. Accordingly, the list comprises several acknowledged repressors including Atf4 [44], Barhl2 [45,46], Bcl11a [47], Foxp1 [48,49], Foxp2 [50], Hey2 [51,52], Scrt1 (= Scratch Family Transcriptional Repressor 1) [53], Six6 [54], Sox2 [55], Tgif1 [56], Tgif2 [56], Vsx1 [57], Vsx2 [58], Zeb2 [59], and Zbtb12 [60]. Of interest, the list of putative repressors comprises three TF that have been labelled as “pioneer factors” (i.e. TF that engage with closed chromatin to open it and make it subsequently accessible to other TF [61]), including Ascl1 [62,63], Sox2 [62], and Isl1 [64]. If the pioneer factor were transiently expressed in retinal progenitor cells (as observed for Ascl1 and Sox2, but not Isl1; STable 12), rendering chromatin accessible to activator TF that are expressed and operate in later stages of development, this may conceivably also generate the observed depletion of binding motifs in the ATAC-Seq peaks of genes that are over-expressed in the same cell type as the pioneer TF. We note that three of the TF in the list of putative repressors (Ahr, Hif1α and Clock), which are typically regarded as paradigmatic activators, are functionally connected: Ahr competes with Hif1α for binding to the nuclear translocator protein Arnt and – possibly - with Clock for binding to Arnt-similar Bmal1 [65]. It is tempting to speculate that this connection may underpin the fact that these three supposedly activator TF appear as repressors in our analyses.

For 10 TF (Etv1, Fosb, Hes1, Jun, Junb, Lhx2, Otx2, Pax6, Sox11 and Tcf3, STable 12), overexpression of the TF was accompanied by binding motif enrichment in some cell type(s), and depletion in other(s). We labelled these as “conditional activator” TF, meaning that the presence of the TF is necessary but not sufficient to exert its effect on transcription. This could for instance reflect the need for post-translational modification of the TF [66], or for cooperation with other TF or cofactors [67], or the fact that the target sites of the TF are not yet accessible requiring further chromatin remodeling [39].

It is increasingly recognized that many TF act both as activator or repressor in different or even in the same cell type, depending (i) on the combination of TF that bind to a given cis-acting regulatory element, as well as (ii) the coregulators (devoid of own DNA binding domain) that they recruit to the regulatory element [67,68]. Accordingly, for many of the TF listed in STable 12, both activator and repressor effects have been reported in the literature. The approach that we have used (i.e. searching for an enrichment or a depletion of binding motifs for an overexpressed TF in the ATAC Seq peaks of all other overexpressed genes) reveals the activity of the TF (activation vs repression) that predominates in a given cell type. For the 31 TF whose predominant activity was of the repressor type, we could nearly always identify one or more cell types in which the ATAC-Seq peaks of over-expressed genes were enriched in the corresponding binding motif (despite the fact that the TF was not over-expressed in that cell type). This strongly suggests that the corresponding repressor specifically targets genes that define another retinal cell type. We revealed a clear relationship between the cell type(s) in which the repressor TF is expressed and the cell type(s) in which its target genes are expressed, allowing us to define three clusters: (I) NE, ERPC, LRPC and NRPC, (II) RGC, AC and HC, and (III) PRP, C, R, BC and MC (Fig. 5D&E). Repressors TF expressed in cluster I (precursor cells) primarily target genes defining cell types of cluster III (primarily photoreceptor and bipolar cells), repressor TF expressed in cluster III primarily target genes defining cell types of cluster I, and repressor TF expressed in cluster II target genes defining cell types of both clusters I and III. Based on these findings, we propose that combinations of activator and repressor TF constitute regulatory toggles that help ensure cell type-specific gene expression and hence cellular identity (Fig. 5C). Our results suggest that the hypothesized regulatory toggles involve multiple activators and repressors. This may confer robustness to the system, and enable differentiation of multiple cell types. It also indicates that perturbing TF one at a time, whether by overexpression or knock-out/down, may not be effective to dissect such multifactorial toggles. It may be necessary to perform pooled screens using CRISPR libraries targeting several candidates at once (at high multiplicity of infection) in Perturb-Seq like experiments conducted in 3D-RA [69] in order to induce detectable alterations in cellular behavior.

We took advantage of publicly available scATAC-Seq data of adult (P56) retina of the mouse (providing information about cell type clusters II and III) [39] that allowed us to test two components of our regulatory toggle hypothesis in an independent data set interrogated with a distinct technology: the effect of 11 TF predicted to operate as activators in cluster III, and the effect of 4 TF predicted to operate as repressors in cluster II. Our analyses provide a vivid visual illustration of (i) the coincident (i.e. in the same cell type) expression of activator TF and their target genes (measured by increased transposase accessibility of both TF-encoding gene and genome-wide ATAC-Seq peaks encompassing the cognate binding motif, respectively), and (ii) the discrepant (i.e. in distinct cell types) expression of repressor TF and their target genes measured in the same manner. Visual impressions were substantiated by highly significant positive (activator TF) and negative (repressor TF) correlations between the transposase accessibility of the TF-gene and the transposase accessibility of ATAC-Seq peaks encompassing the cognate binding-motifs (Fig. 6 and SFig. 9). While the marked contrasting behavior of predicted activator and repressor TF in this assay supports the pertinence of our model, its biological interpretation is not trivial. It is easy to understand that if an activator TF is expressed in a given cell (as testified by the openness of the chromatin surrounding it) and if it is active in that cell, regulatory elements to which it binds (by recognizing cognate motifs in it) to activate target genes will be open and hence accessible as well. But what about the opposite pattern observed for candidate repressor TF? The fact that ATAC-Seq peaks encompassing binding motifs for a repressor are primarily closed in the cell type in which the repressor is expressed suggests that the repressor TF is largely effective and that it contributes to closing the regulatory elements to which it has or is bound. The fact that these same peaks are open in cell types in which the repressor is not expressed suggests that the corresponding regulatory elements encompass binding motifs for both activator and repressor TF. This prediction is substantiated by the observation of strong positive correlations between the density of binding motifs (number of binding motifs divided by peak size) for repressor and activator TF in the utilized scATAC-Seq data (SFig. 11). Thus, our data suggest that the same regulatory elements are used to activate gene expression in the cell type(s) where the gene product is needed, as well as to repress gene expression in the cell type(s) in which expression of the gene is unwanted.

We show that 3D-RA broadly recapitulate the in vivo developmental program and trajectories. However, developmental trajectories appear less canalized in 3D-RA when compared to NaR, PRP to develop earlier and at the expense of other cell types, and terminal differentiation of BC to be incomplete (Fig. 2). We identify ∼3,000 genes that are differentially expressed between 3D-RA and NaR in at least one cell type, and identify the corresponding biological pathways pertaining in particular to the rate of cell division which is reduced in 3D-RA RPCs when compared to NaR, post-transcriptional and translational mechanisms which appear exacerbated in the majority of 3D-RA cell type when compared to NaR, signal transduction via WNT and Notch pathways which are diminished in 3D-RA RPCs when compared to NaR, 3D-RA differentiated cells which appear less functional with less phototransduction cascade activity and decrease synapse formation, and finally apoptosis and stress response which are increased at the latest stages of 3D-RA culture. As for NaR, several of these perturbed pathways were highlighted before in analyses of bulk scRNA-Seq data obtained during the development of NaR and 3D-RA [40], and can now be assigned to cell type-specific transcriptome perturbations. Strikingly, the perturbed pathways show a highly significant overlap with those that were shown to be differentially expressed during the in vivo development of NaR. We show that TF that are differentially expressed during in vivo retinal development are particularly sensitive to the iPSC culture conditions. This is likely to drive the perturbations of the above-mentioned biological pathways.

We show how scRNA-Seq and bulk ATAC-Seq can be combined to gain novel insights into what may underpin differences observed between the NaR and 3D-RA transcriptomes. As an example, the comparison between MC from NaR and 3D-RA revealed 418 genes that were more strongly expressed in NaR when compared to 3D-RA (NaR > 3D-RA), and 424 that were more strongly expressed in 3D-RA (3D-RA > NaR) (STable 7). The first list of genes (NaR > 3D-RA) was not enriched for any Reactome pathway, but the second (3D-RA > NaR) highlighted 53 of these (STable 8), corresponding to five subsets of “found entities” (five colors in MC column (3D-RA > NaR) in Fig. 3B). One of these subsets (orange set, rich in genes encoding ribosomal proteins) highlighted pathways related to RNA stability (NMD), RNA translation, selenocystein metabolism and signaling by ROBO receptors. Combined scRNA-Seq and ATAC-Seq data indicated that genes with high density of binding motifs for Etv1, Etv5, Hes1 and Zbtb7a in their ATAC-Seq peaks are underexpressed in MC of NaR relative to 3D-RA (Fig. 8E&F). These four TF are expressed at relatively high and comparable levels in both NaR and 3D-RA. We identified the corresponding genes and subjected them to Reactome analysis. They identified – to a large extent – the same pathways as the orange gene subset defined above. Thus, the differences between NaR and 3D-RA MC with regards to the corresponding pathways are most likely driven by perturbations of the Etv1, Etv5, Hes1 and Zbtb7a group of TF. Examination of their transcriptional and binding motif enrichment/depletion profile across cell types (Fig. 8D) suggests that Etv1, Etv5 and Hes1 operate as “conditional TF” (as defined above) in NaR: despite still being present in MC they do not have the activator effect in these cells as seen in retinal progenitor cells (hence the observed depletion in the NaR MC transcriptome). In 3D-RA MC, they may still have “residual” activator activity which would explain why the depletion is not seen. Zbtb7a is clearly distinct from the other three: it is most strongly expressed in PRP, C and R, yet its binding motifs are primarily found in genes expressed in retinal progenitor cells. How and why genes enriched in Zbtb7a motifs would be underexpressed in NaR but not 3D-RA MC remains unclear. Yet these examples show how studying the regulatory toggle landscape may become a valuable approach to monitor how closely organoids recapitulate native development.

## Materials and methods

### Generation of iPSC-derived retinal aggregates

#### Maintenance of iPSCs

The mouse iPSC-NrlGFP line was obtained from the laboratory of Retinal Regeneration from the RIKEN Center for Developmental Bioloy (CDB) (Kobe, Japan). These iPSCs were generated from fibroblasts [70] of C57BL/6 Nrl-eGFP transgenic mice [20]. iPSCs were maintained according to [71] in 60-mm Petri dishes (0,6 × 10^5^ cells total per dish) coated with 0.1% gelatin (G2625, Merck, Darmastadt, Germany) in Glasgow’s Minimum Essential Medium (GMEM, 11710035,Thermo Fisher Scientific, Waltham, MA) supplemented with 10% Fetal Bovine Serum (FBS, #04-001-1, Biological Industries, Beit HaEmek, Israel), 1 mM sodium pyruvate (Merck), 0.1 mM MEM Non-Essential Amino Acids Solution (NEAA, Thermo Fisher Scientific), 0.1 mM 2-mercaptoethanol (2-ME, Wako Pure Chemical, Osaka, Japan), 100 U/mL penicillin-streptomycin (Thermo Fisher Scientific), 1000 U/mL of Leukemia inhibitory factor (Esgro LIF, Merck), 3 µM CHIR99021 (BioVision, Milpitas, CA) and 1 µM PD0325901 (Stemgent, Cambridge, MA).

#### Generation of iPSC-derived retinal aggregates

Differentiation of iPSCs into retinal aggregates was done using the SFEBq (serum-free floating culture of embryoid body-like aggregates with quick re-aggregation) method according to [21] with some modifications following [71] and [72]. The iPSCs were dissociated (DD0) after 4-5 days of maintenance using 0.25% trypsin / 1 mM EDTA (Thermo Fisher Scientific) at 37°C for 2 minutes. Embryoid body-like aggregates were formed by adding 5,000 cells/dish in a low binding 96-well microplate (174925, Nunclon Sphera, Thermo Fisher Scientific) in 100 µL of differentiating medium. The differentiating medium is composed of GMEM (Thermo Fisher Scientific), 0.1 mM AGN193109 (Toronto Research Chemicals, Toronto, Canada), 5% of Knock-out Serum Replacement (KSR, Thermo Fisher Scientific), 1 mM Sodium Pyruvate (Merck), 0.1 mM NEAA (Thermo Fisher Scientific) and 0.1 mM 2-ME (Wako). At DD1, 20 µL of Matrigel Growth Factor Reduced Basement Matrix (Corning, Corning, NY) was added to obtain a final concentration equal to 2%. The cells were left in this medium untill DD8. At DD8, retinal aggregates were picked up and transferred in 60-mm Petri dishes in maturation medium composed of Dulbecco’s Modified Eagle’s Medium (DMEM)/F-12 with glutamax (Thermo Fisher Scientific), 1% of N-2 supplement (Thermo Fisher Scientific) and 100 U/mL penicillin-streptomycin (Thermo Fisher Scientific). 0.5 µM retinoic acid (DD13 to DD18) (#R2625, Merck), 1 mM of L-taurine (DD13 to DD29) (#T8691,Merck) and 1% FBS (DD21 to DD29) (Biological Industries) were added to this maturation medium. Taurine and retinoic acid promote rod photoreceptors differentiation [73]. From DD8 to DD29 cultures were maintained in hyperoxic conditions (37°C, 40% O_2_ / 5% CO_2_). Development of retinal aggregates was monitored and GFP expression was confirmed from DD18 using an EVOS FL digital inverted fluorescence microscope (Thermo Fisher Scientific).

### Immunofluorescence

Retinal aggregates were fixed for 20 minutes at room temperature in 4% paraformaldehyde (PFA) in phosphate saline (PBS) at pH 7.4. They were equilibrated overnight in 30% sucrose (in PBS) at 4°C before cryoprotection. Eyeballs from wild type C57BL/6 mice, used as positive controls, were enucleated and punctured in the center of the cornea before fixation for 1 hour in 4% PFA and at room temperature, then washed in PBS and incubated in sucrose 30% at 4°C overnight. Samples were embedded in Richard-Allan Scientific NEG-50 Frozen Section medium (Thermo Fisher Scientific). Slices of 10 to 15 µm were generated with a cryostat and placed on Superfrost Ultra Plus slides (Thermo Fisher Scientific). For immunofluorescence, slides were first incubated in Blocking One solution (Nacalai Tesque, Kyoto, Japan) for 1 hour at room temperature, then at 4°C overnight with primary antibodies diluted in Dako REAL Antibody Diluent (Agilent, Santa Clara, CA). We used the following primary antibodies: rabbit antibody against Protein Kinase Cα diluted at 1:500 (Antibody Registry ID: AB_477345, Merck), rabbit antibody against Recoverin at 1:1000 (AB_2253622, Merck), rabbit antibody against Calretinin at 1:500 (AB_2313763, Swant, Marly, Switzerland), rabbit antibody against Pax6 at 1:100 (AB_2313780, BioLegend, San Diego, CA), mouse antibody against RET-P1 at 1:1000 (anti-Rhodopsin, AB_260838, Merck), sheep antibody against Chx10 at 1:1000 (AB_2314191, Exalpha Biologicals, Shirley, MA). After 24 hours, slides were washed three times for 5 minutes in 0.05% PBS-Tween then incubated with appropriate secondary antibodies in the dark at room temperature (anti-IgG rabbit A488 and A647, anti-IgG mouse A555 and anti-IgG sheep A555, all from Thermo Fisher Scientific) and 1:1000 4’,6-diamidino-2-phenylindole (DAPI) in Dako REAL Antibody Diluent. After another wash in PBS-Tween, slides were mounted with FluorSave Reagent (Merck). Images were taken with a Nikon Eclipse T_i_ confocal microscope.

### Single cell RNA Seq

#### Dissociation of native retinal tissue and 3D-culture retinal aggregates

The dissociation of mouse retinas and 3D retinal aggregates was inspired by the protocol of Macosko et al. [74]. Eyeballs of C57BL/6 wild type mice were enucleated at time points E13, P0, P5 and P9. Dissected retinas were placed in Dulbecco’s Phosphate Buffered Saline (DPBS, Thermo Fisher Scientific). Optic vesicule (OV)-like structures of the iPSCs derived 3D retinal aggregates were cut at DD13, DD21, DD25 and DD29 and transferred in DPBS as well. Papain 4 U/mL (Worthington Biochemical Corporation) was added to the samples. The solution containing the retinas and the OV-like structures was maintained at 37°C for 45 and 30 minutes, respectively. 0.15% ovomucoid (Worthington Biochemical Corporation, Lakewood, NJ) was added for papain inhibition. Samples were centrifuged in order to eliminate the supernatant and cells were resuspended in DPBS. Cell numbers and proportion of living cells were estimated by Trypan Blue staining and using a Countess II cell counter (Thermo Fisher).

#### scRNA-Seq

We generated two biological replicates for stages 1 to 3 (NaR and 3D-RA) and one biological replicate for stage 4 (NaR and 3D-RA). We loaded ∼15,700 cells for biological replicate 1 (stage 1-4) and ∼10,000 cells for biological replicate 2 (stage 1-3) in a Chromium Controller instrument (10X Genomics, Pleasanton, CA). Sequencing libraries were generated using Chromium Single Cell 3’ reagent kits v2.0 following the recommendations of the manufacturer (10X Genomics). Actual sequencing was conducted on an Illumina NextSeq 500 instrument (Illumina, Sand Diego, CA).

#### Bioinformatic analyses

Demultiplexing, alignment, filtering, barcode counting, UMI counting, and aggregation of multiple runs were conducted using Cell Ranger v2.1.1 (10X Genomics). Further filtering, k-means clustering, UMAP projection were conducted using the Seurat software suite (https://satijalab.org/seurat/)[23]. Velocity analysis was performed using the Velocyto R [11] and scvelo [12] packages. Single-cell trajectory inference and pseudotime analyses were conducted using Monocle2 (http://cole-trapnell-lab.github.io/monocle-release/)[14].

### ATAC-Seq

#### Data generation

ATAC-seq libraries were constructed on NaR (E13, P0, P5) and 3D-RA (DD13, DD21, DD25) samples with biological replicates following the Omni ATAC protocol [36]. We used 50,000 cells per reaction taken from the cell suspensions prepared for the scRNA-seq. We tested two different amounts of Tagment DNA TDE1 enzyme (1 and 2 μl in a 50 μl reaction volume) (Illumina) per sample. Genomic DNA (gDNA) libraries were also prepared using 50 ng of gDNA isolated from NaR P5 and 3D-RA DD25 cells by following the Nextera DNA Sample Preparation Guide (Illumina). The libraries were purified using the MinElute PCR purification kit (Qiagen, Venlo, Netherlands) followed by 13 and 5 cycles of PCR-amplifications for ATAC-seq and gDNA libraries, respectively. After validating library size distribution using the QIAxcel capillary electrophoresis (Qiagen), the libraries were further purified using the SPRIselect reagent to remove large DNA molecules (a right-side size selection with 0.55X followed by 1.5X ratios of beads) (Beckman Coulter, Brea, California). On average 10.6 millions of 38-nucleotide paired-end sequences were obtained using a NextSeq 500 sequencer (Illumina).

#### Data analyses

Data was analyzed by following the ENCODE Kundaje lab ATAC-seq pipeline (https://www.encodeproject.org/pipelines/ENCPL792NWO/). Sequences were trimmed using Trimmomatic [75] and aligned on the Mus musculus genome assembly mm10 using Bowtie2 [76]. After filtering out low quality, multiple mapped, mitochondrial, and duplicated reads using SAMtools [77] and the Picard Toolkit (http://broadinstitute.github.io/picard/), fragments with map length ≤ 146 bp were kept as nucleosome-free fraction. Genomic loci targeted by TDE1 were defined as 38-bp regions centered either 4 (plus strand reads) or 5-bp (negative strand reads) downstream of the read’s 5’-end. ATAC-seq peaks were called using the MACS2 software (narrowPeak; q-value ≤ 0.01) [38]. FRiP scores were calculated as the fraction of TDE1 targeted loci falling into the called peaks. Overlapping peaks across samples were merged and annotated for the occurrence of TF binding motifs of interest (Suppl. Table 11) and the closest gene using Homer [38]. TDE1 targeted loci overlapping the merged peaks were extracted and converted to a bedgraph file with a scaling factor to one million reads using BEDTools [78], and further to tdf format to visualize peaks on the Integrative Genomics Viewer [79]. The total number of TDE1 targeted loci overlapping the merged peaks were counted using BEDOPS [80], normalized for peak lengths and a sequencing depth with per one million scaling factor, standardized and used for hierarchical cluster analysis using R hclust [81] and gplots (https://CRAN.R-project.org/package=gplots). The detailed analysis pipeline is provided in the ATAC_seq_analysis_pipeline.docx file. The overall mapping rate with Bowtie2 averaged 98.6%, the mapping rate to the mitochondrial genome 4.1%, the duplicate fragment rate 6.0%, the proportion of usable reads after filtration 83.4%, and the FRiP score 34.1%. The FRiP score was significantly lower for E13 samples (reminiscent of the E14.5 samples in [82]), yet not so in the equivalent DD13 samples (Suppl. Table 16).

### Accessing publicly available scATAC-Seq and bulk ATAC-seq

Single-cell ATAC-seq data from 8-wk wild-type C57BL/6 mouse retinas were obtained from GEO:GSE164044 [39] and analyzed using Cell Ranger ATAC v1.2.0 and Loupe Browser v5.0 with default settings (10X Genomics). Bulk ATAC-seq data on FACS-sorted rod and cone photoreceptors from 8-wk Nrl-eGFP and Opn1mw-GFP mouse, respectively, were obtained from GEO:GSE83312 [83] and analyzed as above.

### Downstream analyses

#### Width of developmental trajectories in 2D UMPA space

To test whether the developmental trajectories were more tightly regulated in NaR than in 3D-RA we computed the average distance (computed as the Euclidian distance in 2D-UMAP space, i.e.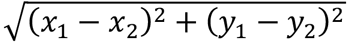 between 500 randomly selected NaR and 500 randomly selected 3D-RA cells and their *n* nearest neighbors (with *n* ranging from 1 to 50). The number of cells per developmental stage was adjusted between NaR and 3D-RA by down sampling to the number of the least populated source. The corresponding calculations were performed five times. The curves shown in Fig. 2D correspond to the averages across the five replicates. The grey confidence zone in Fig. 2D is bounded by the maxima and minima across the five replicates. The corresponding script was written in Perl (Dev_path_width.pl) and the graph generated in R (Path_width.R).

#### Within developmental stage cell type entropy

To compare cell type diversity within developmental stage between NaR and 3D-RA, we first equalized the number of cells with developmental stage between NaR and 3D-RA by randomly dropping cells from the most populated source. We then sampled two cells within cell source (NaR and 3D-RA) and developmental stage and checked whether they were from the same cell type or not. This was repeated 1,000 times yielding a measure of cell type diversity akin to (1-entropy). Down-sampling of cells was repeated 100 times. Each data point in Fig. 2E corresponds to (1-Entropy) for one such random sample. The corresponding script was written in Perl (entropy.pl) and the graph generated in R (Entropy.R).

#### Differential expression analyses

Differential expression analyses to identify genes that are upregulated in specific cell types when compared to all other ones (Cell type > Others) or that are differentially expressed between NaR and 3D-RA in a given cell type (Nar > 3D-RA and 3D-RA > NaR) were performed with the *Findmarkers* function in Seurat (https://satijalab.org/seurat/). *<u>Pathway analyses:</u>* Pathway enrichment analyses were conducted using the on-line Reactome analysis tools [33,34]. Mouse gene identifiers were converted to human counterparts. Pathway analysis results were downloaded as flat files. A total of 392 pathways with enrichment p-value ≤ 0.01 in at least one analysis were kept and manually sorted according to Reactome hierarchy (Man_processed_reactome_output.txt). A pathway is enriched in a list of genes if it contains more components of the pathway than expected by chance (given the number of genes in the list). The overlapping genes (“Found entities”) hence define the enrichment. The same pathway can be enriched in two gene lists due to the same, distinct or partially overlapping sets of “found entities”. We quantified the degree of overlap between sets of “found entities” for the 1,313 pathway enrichments using principal component (PC) analysis in a space defined by the presence/absence of 1,335 genes. The distance between sets of “found entities” in a space consisting of the 20 first PCs was projected in 3D space using t-distributed stochastic neighbor embedding (tSNE) implemented with the *Rtsne* R function. 3D tSNE coordinates were converted to hexadecimal RGB code and used to color the sets of “found entities” (corresponding to the enrichment of a pathway in a specific gene list) when generating 2D tSNE graphs (SFig. 4), or when generating a tile showing the pathways enriched in specific analyses (Cell type>OTHER, NaR > 3D-RA or 3D-RA > NaR) and cell type within analysis (NE, RPE, ERPC, LRPC, NRPC, RGC, HC, AC, PRP, C, R, BC or MC) (Fig. 3B). The corresponding scripts were written in Perl (Reactome_analysis.pl) and R (Reactome_analysis.R).

#### Identifying regulatory toggles

We used Homer [38] to compile the number of occurrences of 336 binding motifs for 151 of 307 dynamically regulated TF in 98,181 ATAC-Seq peaks assigned to 19,170 genes. For each gene, the data were summarized as (i) the <u>total</u> number of occurrences, and (ii) the <u>mean</u> number of occurrences per peak (i.e. density), for each of the 336 binding motifs (STable 11). We then checked - for each of the 336 binding motifs separately - whether the number (“total” in STable 11) and density (“mean” in STable 11) of motifs differed significantly between genes that were upregulated versus downregulated in every one of the 13 cell types. Differential expression analyses to identify genes that are up- and downregulated in specific cell types were performed with the *Findmarkers* function in Seurat (https://satijalab.org/seurat/). The corresponding results are summarized in a series of files labelled, respectively, “NaR/RET_<CELL_TYPE>_markers.txt” for NaR, and “3D_RA/IPS_<CELL_TYPE>_markers.txt” for 3D_RA. We used a threshold q-value of 0.05 to declare a gene as significantly up- or down-regulated in a given cell type. The statistical significance of the difference in number and density of binding motifs between up- and down-regulated genes was computed using Wilcoxon rank-based test implemented with the wilcox.test R function. Differences were deemed significant if the q-value (computed with the qvalue R function) was ≤ 0.01. Corresponding results are provided as STable 13 for NaR and STable 14 for 3D-RA. The graphs for figure 5 were generated using the Comb_scRNA_ATAC_seq R script.

All used scripts and datasets are available without restrictions from: http://web.giga.ulg.ac.be/pubdata/UAG/Georges_A_2020.

## AUTHOR CONTRIBUTIONS

Conceived and designed the experiments: AG, MT, MM, HT, MG. Performed the experiments: AG, HT, FL, LK, SD. Analyzed the data: AG, AL, HT, MS, LD, MG. Contributed reagents/materials/analysis tools/supervision: AG, AL, LN, JMR, LD, MS, MT, MG. Wrote the paper: AG, HT, MG.

## ACKNOWLEDGEMENTS

We are grateful to Tomoyo Hashiguchi for teaching the 3D-RA culture protocol and to Tomohiro Masuda and Akishi Onishi for their comments and suggestions. We thank the GIGA Genomics platform for their help both with scRNA-seq experiments and early bio-informatic analyses (especially Wouter Coppieters and Benoit Charloteaux). We are grateful to Vincent Lambert for assisting with the dissection of NaR. This work was financially supported by the “King Baudouin Foundation”, the “Global Ophthalmology Awards Program (GOAP) from Bayer”, and the “Fonds Leon Fredericq”. LD and LN are respectively postdoctoral researcher and senior associate researcher of the FRS-FNRS. We are grateful to the Dyer laboratory for the scATAC-Seq data.

## DATA AVAILABILITY

All data generated as part of this work are available without restrictions. They have been deposited under accession numbers E-MTAB-9440 and E-MTAB-9395. All data and analysis pipelines are available at http://web.giga.ulg.ac.be/pubdata/UAG/Georges_A_2020. The authors declare that they have no competing interests.

## ETHICAL APPROVAL

Ethical approval: All animal procedures were approved by the Animal Ethics Committee at University of Liège (approval no. 17-1908) and performed in accordance with the Guide for the Care and Use of Laboratory Animals at University of Liège.

## REFERENCES

1. Eiraku M, Takata N, Ishibashi H, Kawada M, Sakakura E, Okuda S, Sekiguchi K, Adachi T, Sasai Y. Self-organizing optic-cup morphogenesis in three-dimentional culture. Nature. 2011; 472:51–56.

2. Meyer JS, Howden SE, Wallace KA, Verhoeven AD, Wright LS, Capowski EE, Pinilla I, Martin JM, Tian S, Stewart R, et al. Optic vesicle-like structures derived from human pluripotent stem cells facilitate a customized approach to retinal disease treatment. Stem Cells. 2011; 29:1206–1218.

3. Nakano T, Ando S, Takata N, Kawada M, Muguruma K, Sekiguchi K, Saito K, Yonemura S, Eiraku M, Sasai Y. Self-formation of optic cups and storable stratified neural retina from human ES cells. Cell Stem Cell. 2012;10:771–785.

4. Li JQ, Welchowski T, Schmid M, Letow J, Wolpers AC, Holz FG, Finger RP. Retinal diseases in Europe. Euretina Whitebook 2017; https://www.euretina.org/downloads/EURETINA_Retinal_Diseases.pdf.

5. Jin ZB, Okamoto S, Osakada F, Homma K, Assawachananont J, Hirami Y, Iwata T, Takahashi M. Modeling retinal degeneration using patient-specific induced pluripotent stem cells. PLoS One. 2011;6:e17084.

6. Achberger K, Haderspeck JC, Kleger A, Liebau S. Stem cell-based retina models. Advanced Drug Delivary Reviews. 2019; 140:33–50.

7. Völkner M, Zschätzsch M, Rostovskaya M, Overall RW, Busskamp V, Anastassiadis K, Karl MO. Retinal organoid from pluripotent stem cells efficiently recapitulate retinogenesis. Stem Cells Rep. 2016;6:525–538.

8. Shekhar K, Lapan SW, Whitney IE, Tran NM, Macosko EZ, Kowalczyk M, Adiconis X, Levin JZ, Nemesh J, Goldman M, McCarroll SA, Cepko CL, Regev A, Sanes JR. Comprehensive classification of retinal bipolar neurons by single-cell transcriptomics. Cell. 2016;166:1308–1323.

9. Saelens W, Cannoodt R, Todorov H & Saeys Y. A comparison of single cell trajectory inference methods. Nat Biotechnol. 2019;37:547–554.

10. Haghverdi L, Büttner M, Wolf FA, Buettner F, Theis FJ. Diffusion pseudotime robustly reconstructs lineage branching. Nat Methods 2016;13:845–848.

11. La Manno G, Soldatov R, Zeisel A, Braun E, Hochgerner H, Petukhov V, Lidschreiber K, Kastriti ME, Lonnerberg P, Furlan A, Fan J, Borm LE, Liu Z, Van Bruggen D, Guo J, He X, Barker R, Sundstrom E, Castelo-Branco G, Cramer P, Adameyko I, Linnarsson S, Kharchenko P. RNA velocity of single cells. Nature. 2018;560:494–510.

12. Bergen V, Lange M, Peidli S, Wolf FA, Theis FJ. Generalizing RNA velocity to transient cell states through dynamic modeling. Nat Biotechnol. 2020; 38:1408–1414.

13. Camara PG. Methods and challenges in the analysis of single-cell RNA-sequencing data. Curr Opin Syst Biol. 2018; 7:47–53.

14. Trapnell C, Cacchiarelli D, Grimsby J, Pokharel P, Li S, Morse M, Lennon NJ, Livak KJ, Mikkelsen TS, Rinn JL. The dynamics and regulators of cell fate decisions are revealed by pseudotemporal ordering of single cells. Nat Biotechnol. 2014;32:381–386.

15. Collin J, Queen R, Zerti D, Dorgau B, Hussain R, Coxhead J, Cockell S, Lako M. Deconstructing retinal organoids: single cell RNA-Seq reveals the cellular components of human pluripotent stem cell-derived retina. Stem Cells. 2019;37:593–598.

16. Sridhar A, Hoshino A, Finkbeiner CR, Chitsazan A, Dai L, Haugan AK, Eschenbacher KM, Jackson DL, Trapnell C, Bermingham-McDonogh O, Glass I, Reh TA. Single Cell Transcriptomic Comparison of human fetal retina, hPSC-derived retinal organoids, and long-term retinal cultures. Cell Rep. 2020;30:1644–1659.

17. Cowan CS, Renner M, De Gennaro M, Gross-Scherf B, Goldblum D, Hou Y, Munz M, Rodrigues TM, Krol J, Szikra T, Cuttat R, Waldt A, Papasaikas P, Diggelmann R, Patino-Alvarez CP, Galliker P, Spirig SE, Pavlinic D, Gerber-Hollbach N, Schuierer S, Srdanovic A, Balogh M, Panero R, Kusnyerik A, Szabo A, Stadler MB, Orgül S, Picelli S, Hasler PW, Hierlemann A, Scholl HPN, Roma G, Nigsch F, Roska B. Cell types of the human retina and its organoids at single-cell resolution. Cell. 2020;182:1623–1640.

18. Yan F, Powell DR, Curtis D & Wong NC. From reads to insight: a hitchhiker’s guide to ATAC-Seq data analysis. Genome Biol. 2020;21:22.

19. Gonzalez-Cordero A, West EL, Pearson RA, Duran Y, Carvalho LS, Chu CJ, Naeem A, Blackford SJ, Georgiadis A, Lakowski J, Hubank M, Smith AJ, Bainbridge JWB, Sowden JC, Ali RR. Photoreceptor precursors derived from three-dimensional embryonic stem cell cultures integrate and mature within adult degenerate retina. Nat Biotechnol. 2013;31:741–747.

20. Akimoto M, Cheng H, Zhu D, Brzezinski JA, Khanna R, Filippova E C T, Oh E, Jing Y, Linares JL, Brooks M, Zareparsi S, Mears AJ, Hero A, Glaser T, Swaroop A. Targeting of GFP to newborn rods by Nrl promoter and temporal expression profiling of flow-sorted photoreceptors. Proc Natl Acad Sci USA. 2006;103:3890–3895.

21. Eiraku M, Sasai Y. Mouse embryonic stem cell culture for generation of three-dimensional retinal and cortical tissues. Nat Prot. 2012;7:69–79.

22. Assawachananont J, Mandai M, Okamoto S, Yamada C, Eiraku M, Yonemura S, Sasai Y, Takahashi M. Transplantation of embryonic and induced pluripotent stem cell-derived 3D retinal sheets into retinal degenerative mice. Stem Cell Rep. 2014;2:662–674.

23. Butler A, Hoffman P, Smibert P, Papalexi E, Satija R. Integrating single-cell transcriptomic data across different conditions, technologies, and species. Nat Biotech. 2018;36:411–420.

24. McInnes L, Healy J, Melville J. UMAP: Uniform Manifold Approximation and Projection. J. Open Source Software. 2018;3:861.

25. Clark BS, Stein-O’-Brien GL, Shiau F, Cannon GH, Davis-Marcisak E, Sherman T, Santiago CP, Hoang TV, Rajaii F, James-Esposito RE, Gronostajski RM, Fertig EJ, Goff LA, Blackshaw S. Comprehensive analysis of retinal development at single cell resolution identifies NFI factors as essential for mitotic exit and specification of late-born cells. Neuron. 2019;102:1111–1126.

26. Tiroshi I, Izar B, Prakadan SM, Wadsworth MH, Treacy D, Trombetta JJ, Rotem A, C Rodman, Lian C, Murphy G, Fallahi-Sichani M,Dutton-Regester K, Lin JR, Cohen O, Shah P, Lu D, Genshaft A, Hughes TK, Ziegler CGK, Kazer SW, Gaillard A, Kolb KE, Villani AC, Johannessen CM, Andreev AY, Van Allen EM, Bertagnolli M, Sorger PK, Sullivan RJ,Flaherty RT,Frederick DT, Jané-Valbuena J, Yoon CH, Rozenblatt-Rosen O, Shalek AK, Regev A, Garraway LA. Dissecting the multicellular ecosystem of metatastic melanoma by single-cell RNA-seq. Science. 2016;352:189–196.

27. Trimarchi JM, Cho SH, Cepko CL. Identification of genes expressed preferentially in the developing peripheral margin of the optic cup. Dev Dyn. 2009;238:2327.

28. Liu J, Reggiani JDS, Laboulaye ME, Pandey S, Chen B, Rubenstein JLR, Krishnaswamy A, Sanes JR. Tbr1 instructs laminar pattering of retinal ganglion cell dendrites. Nat Neurosci. 2018;21:659–670.

29. Yan W, Laboulaye MA, Tran NM, Whitney IE, Benhar I, Sanes JR. Mouse retinal cell atlas: molecular identification of over sixty amacrine cell types. J. Neurosci. 2020;40:5177–5195.

30. Bassett EA, Wallace VA. Cell fate determination in the vertebrate retina. Trends in Neurosciences. 2012; 35:565–573.

31. Reese B. Development of the retina and optic pathway. Vision Res. 2011;51:613–632.

32. Kuwahara A, Ozone C, Nakano T, Saito K, Eiraku M, Sasai Y. Generation of a ciliary margin-like stem cell niche from self-organizing human retinal tissue. Nat Commun. 2015;6:6286.

33. Fabregat A, Jupe S, Matthews L, Sidiropoulos K, Gillespie M, Garapati P, Haw R, Jassal B, Korninger F, May B, Milacic M, Roca CD, Rothfels K, Sevilla C, Shamovsky V, Shorser S, Varusai T, Viteri G, Weiser J, Wu G, Stein L, Hermjakob H, D’Eustachio P. The reactome pathway knowledgebase. Nucl Ac Res. 2018;44:D481–487.

34. Jassal B, Matthews L, Viteri G, Gong C, Lorente P, Fabregat A, Sidiropoulos K, Cook J, Gillespie M, Haw R, Loney F, May B, Milacic M, Rothfels K, Sevilla C, Shamovsky V, Shorser S, Varusai T, Weiser J, Wu G, Stein L, Hermjakob H, D’Eustachio P. The reactome pathway knowledgebase. Nucleic Acids Res. 2020;48:D498–D503.

35. Kanamori M, Konno H, Osato N, Kawai J, Hayashizaki Y, Suzuki H. A genome-wide and nonredundant mouse transcription factor database. Biochem Biophys Res Commun. 2004;322:787–793.

36. Corces MR, Trevino AE, Hamilton EG, Greenside PG, Sinnott-Armstrong NA, Vesuna S, Satpathy AT, Rubin AJ, Montine KS, Wu B, Kathiria A, Cho SW, Mumbach MR, Carter AC, Kasowski M, Orloff LA, Risca VI, Kundaje A, Khavari PA, Montine TJ, Greenleaf WJ & Chang HY. An improved ATAC-seq protocol reduces background and enables interrogation of frozen tissues. Nat Methods. 2017;14:959–962.

37. Zhang Y, Liu T, Meyer CA, Eeckhoute J, Johnson DS, Bernstein BE, Nusbaum C, Myers RM, Brown M, Li W & Liu XS. Model-based analysis of ChIP-Seq (MACS). Genome Biol. 2008;9:R137.

38. Heinz S, Benner C, span N, Bertolino E, Lin YC, Laslo P, cheng JX, Murre C, Singh H, Glass CK. Simple combinations of lineage-determining transcription factors prime cis-regulatory elements required for macrophage and B cell identities. Mol Cell. 2010; 38:576–589.

39. Norrie JL, Lupo MS, Xu B, Al Diri I, Valentine M, Putnam D, Griffiths L, Zhang J, Johnson D, Easton J, Shao Y, Honnell V, Frase S, Miller S, Stewart V, Zhou X, Chen X, Dyer MA. Nucleosome dynamics during retinal development. Neuron. 2019; 104:512–528.

40. Brooks MJ, Chen HY, Kelley RA, Mondal AK, Nagashima K, De Val N, Li T, Chaitankar V, Swaroop A. Improved retinal organoid differentiation by modulating signaling pathways revealed by comparative transcriptome analyses with development in vivo. Stem Cell Reports. 2019; 13:891–905.

41. Swaroop A, Kim D, Forrest D. Transcriptional regulation of photoreceptor development and homeostasis in the mammalian retina. Nat Rev Neurosc. 2010; 11:563–576.

42. Brzezinski JA, Reh TA. Photoreceptor cell fate specification in vertebrates. Development. 2015; 142:3263–3273.

43. Buono L, Martinze-Morales J-R. Retina development in vertebrates: systems biology approaches to understanding genetic programs. Bioessays. 2020; 42:e1900187.

44. Smith SG, Haynes KA, Hegde AN. Degradation of transcriptional repressor ATF4 during long-term synaptic plasticity. Int J Mol Sci. 2020; 21:8543.

45. Sena E, Rocques N, Borday C, Amin HSM,Parain K, Sitbon D, Chesneau A, Durand BC. Barhl2 maintains T cell factors as repressors and thereby switches off the Wnt/β-catenin response driving Spemann organizer formation. Development. 2019; 146:dev.173112.

46. Reig G, Cabrejos ME, Concha ML. Functions of BarH transcription factors during embryonic development. Dev Biol. 2007; 302:367–375.

47. Bauer DE, Orkin SH. Hemoglobin switching’s surprise: the versatile transcription factor BCL11A is a master repressor of fetal hemoglobin. Curr Opin Genet Dev. 2015; 33:62–70.

48. https://www.uniprot.org/uniprot/Q9H334

49. Sollis E, Graham SA, Vino A, Froehlich H, Vreeburg M, Dimitropoulou D, Gilissen C, Pfundt R, Rappold GA, Brunner HG, Deriziotis P, Fisher SE. Identification and functional characterization of de novo FOXP1 variants provides novel insights into the etiology of neurodevelopmental disorder. Hum Mol Genet. 2016; 25:546–557.

50. https://www.uniprot.org/uniprot/O15409

51. https://www.uniprot.org/uniprot/Q9UBP5

52. Nakagawa O, McFadden DG, Nakagawa M, Yanagisawa H, Hu T, Srivastava D, Olsen EN. Members of the HRT family of basic helix-loop-helix proteins act as transcriptional repressors downstream of Notch signaling. Proc Natl Acad Sci USA. 2000; 97:13655–13660.

53. Nakakura EK, Watkins DN, Schuebel KE, Sriuanpong V, Borges MW, Nelkin BD, Ball DW. Mammalian Scratch: a neural-specific Snail family transcriptional repressor. Proc Natl Acad Sci USA. 2001; 98:4010–4015.

54. Li X, Perissi V, Liu F, Rose DW, Rosenfeld MG. Tissue-specific regulation of retinal and pituitary precursor cell proliferation. Science. 2002; 297:1180–1183.

55. Liu Y-R, Laghari ZA, Novoa CA, Hughes J, Webster JRM, Goodwin PE, Wheatley SP, Scotting PJ. Sox2 acts as a transcriptional repressor in neural stem cells. BMC Neurosci. 2014; 15:95.

56. Melhuish TA, Taniguchi K, Wotton D. Tgif1 and Tgif2 regulate axial patterning in mouse. PLoS ONE. 2016; 11:e0155837.

57. Dorval KM, Bobechko BP, Ahmad KF, Bremmer R. Transcriptional activity of the paired-like homeodomain proteins CHX10 and VSX1. J Biol Chem. 2005; 280:10100–10108.

58. Vitorino M, Jusuf PR, Maurus D, Kimura Y, Higashijima S-I, Harris WA. Vsx2 in the zebrafish retina: restricted lineages through derepression. Neural Dev. 2009; 4:14.

59. https://www.uniprot.org/uniprot/O60315

60. Siggs OM, Beutler B. the BTB-ZF transcription factors. Cell Cycle. 2012; 11:3358–3369.

61. Zaret KS, Carroll JS. Pioneer transcription factors: establishing competence for gene expression. Genes & Dev. 2011; 25:2227–2241.

62. Morris SA. Direct lineage reprogramming via pioneer factors; a detour through developmental gene regulatory networks. Development. 2016; 143:2696–2705.

63. Raposo AASF, Vasconcelos FF, Drechsel D, Marie C, Johnston C, Dolle D, Bithell A, Gillotin S, van den Berg DLC, Ettwiller L, Flicek P, Crawford GE, Parras CM, Berninger B, Buckley NJ, Guillemot F, Castro DS. Ascl1 coordinately regulates gene expression and the chromatin landscape during neurogenesis. Cell Reports. 2015; 10:1544–1556.

64. Gao R, Liang X, Cheedipudi S, Cordero J, Jiang X, Zhang Q, Caputo L, Günther S, Kuenne C, Ren Y, Bhattacharya S, Yuan X, Barreto G, Chen Y, Braun T, Evans SM, Sun Y, Dobreva G. Pioneering function of Isl1 in the epigenetic control of cardiomyocyte fate. Cell Research. 2019; 29:486–501.

65. Gargaro M, Scalisi G, Manni G, Mondanelli G, Grohmann U, Fallarino F. The landscape of Ahr regulators and coregulators to fine-tune Ahr functions. Int J Mol Sci. 2021; 22:757

66. Filtz TM, Vogel WK, Leid M. Regulation of transcription factor activity by interconnected, post-translational modifications. Trends Pharmacol Sci. 2014;35:76–85.

67. Lambert SA, Jolma A, Campitelli LF, Das PK, Yin Y, Albu M, Chen X, Taipale J, Hughes TR, Weirauch MT. The human transcription factors. Cell. 2018;172:650–665.

68. White MA, Kwasnieski JC, Myers CA, Shen SQ, Corbo JC, Cohen BA. A simple grammar defines activating and repressing cis-regulatory elements in photoreceptors. Cell Reports. 2016; 17:1247–1254.

69. Dixit A, Parnas O, Li B, Fulco CP, Jerby-Arnon L, Marjanovic ND, Dionne D, Burks T, Raychowdhury R, Adamson B, Norman TM, Lander ES, Weissman JS, Friedman N, Regev A. Perturb-Seq: dissecting molecular circuits with scalable single-cell RNA profiling of pooled genetic screens. Cell. 2016;167:1853–1866.

70. Homma K, Okamoto S, Mandai M, Gotoh N, Rajasimha HK, Chang YS, Chen S, Cogliati T, Swaroop A, Takahashi M. Developing rods transplanted into the degenerating retina of Crx-knockout mice exhibit neural activity similar to native photoreceptors. Stem Cells. 2013;31:1149–1159.

71. Iwasaki Y, Sugita S, Mandai M, Yonemura S, Onishi A, Ito SI, Mochizuki M, Ohno-Matsui K, Takahashi M. Differentiation/purification protocol for retinal pigment epithelium from mouse induced pluripotent stem cells as a research tool. PLoS One. 2016;11:1–20.

72. Eiraku M, Takata N, Ishibashi H, Kawada M, Sakakura E, Okuda S, Sekiguchi K, Adachi T & Sasai Y. Self-organizing optic-cup morphogenesis in 3D culture. Nature. 2011;472:51–56.

73. Osakada F, Ikeda H, Mandai M, Wataya T, Watanabe K, Yoshimura N, Akaike A, Sasai Y & Takahashi M. Toward the generation of rod and cone photoreceptors from mouse, monkey and human embryonic stem cells. Nat Biotechnol. 2008;26:215–224.

74. Macosko EZ, Basu A, Satija R, Nemesh J, Shekhar K, Goldman M, Tirosh I, Bialas AR, Kamitaki N, Martersteck Trombetta JJ, Weitz DA, Sanes JR, Shalek AK, Regev A, McCarroll SA. Highly parallel genome wide expression profiling of individual cells using nanoliter droplets. Cell. 2015;161:1202:1214.

75. Bolger AM, Lohse M, Usadel B. Trimmomatic: a flexible trimmer for Illumina sequence data. Bioinformatics. 2014; 30:2114–2120.

76. Langmead A, Salzberg S. Fast gapped-read alignment with Bowtie 2. Nature Methods. 2012 ; 9 : 357–359.

77. Li H, Handsaker B, Wysoker A, Fennell T, Ruan J, Homer N, Marth G, Abecasis G, Durbin R. 1000 Genome Project Data Processing Subgroup. The Sequence Alignment/Map format and SAMtools. Bioinformatics. 2009;25:2078–2079.

78. Quinlan AR & Hall IM. BEDTools: a flexible suite of utilities for comparing genomic features. Bioinformatics. 2010;26:841–842.

79. Robinson JT, Thorvaldsdottir H, Winckler W, Guttman M, Lander ES, Getz G & Mesirov JP. Integrative genomics viewer. Nat Biotechnol. 2011;29:24–26.

80. Neph S, Kuehn MS, Reynolds AP, Haugen E, Thurman RE, Johnson AK, Rynes E, Maurano MT, Vierstra J, Thomas S, Sandstrom R, Humbert R, Stamatoyannopoulos JA. BEDOPS: high performance genomic feature operations. Bioinformatics. 2012;28:1919–1920.

81. Murtagh F, Legendre P. Ward’s hierarchical agglomerative clustering method: which algorithms implement Ward’s criterion ? J Classification. 2014;31:274–295.

82. Aldiri I, Xi B, Wang L, Chan X, Hiler D, Griggiths L, Valentine M, Shirinifard A, Thiagarajan S, Sablauer A, Barabas M-E, Zhang J, Johnson D, Frase S, Zhou X, Easton J, Zhang J, Maris ER, Wilson RK, Downing JR, Dyer MA. The dynamic epigenetic landscape of the retine during development, reprogramming and tumorigenesis. Neuron. 2017. 94: 550–568.

83. Hughes AE, Enright JM, Myers CA, Shen SQ, et al. Cell type-specific epigenomic analysis reveals a uniquely closed chromatin architecture in mouse rod photoreceptors. Sci Rep. 2017; 7:43184.

