## Supplemental Material for "Evidence from combined analysis of single cell RNA-Seq and ATAC-Seq data of regulatory toggles operating in native and iPS-derived murine retina"

[1]GIGA Stem Cells, GIGA Institute, University of Liège, Belgium. [2]Department of Ophthalmology, Faculty of Medicine and CHU University Hospital, University of Liège, Belgium. [3]GIGA Medical Genomics, GIGA Institute, University of Liège, Belgium. [4]GIGA Genomics Platform, GIGA Institute, University of Liège, Belgium. [5]Laboratory for Retinal Regeneration, Center for Developmental Biology, RIKEN, Japan. [6]Digital Business, HEC Management School, University of Liège, Belgium. [7]Senior Visiting Scientist, Laboratory for Retinal Regeneration, Center for Biosystems Dynamics Research, RIKEN, Japan.

### Supplemental Figure 1:

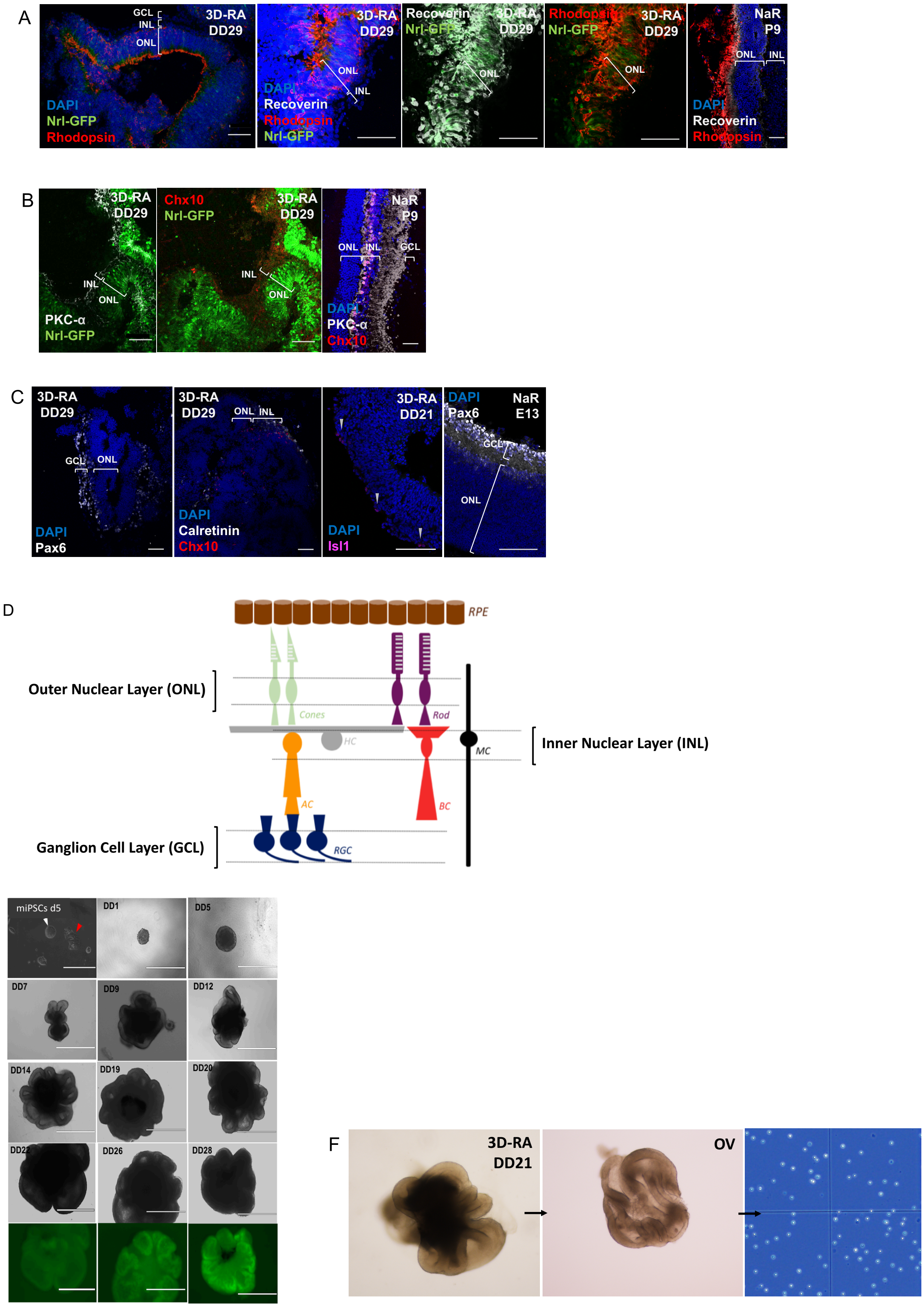

**(A - C) Expected layered expression of cell-type specific immunohistochemical markers.** Immunohistochemical markers of cellular subtypes in NaR (E13 and P9) and Nrl-GFP iPSCs-derived 3D-RA (DD21 and DD29). **(A)** Photoreceptor cells**:** Nrl-GFP (green), Recoverin (white) and Rhodopsin (red) are specific for rod photoreceptor cells; DAPI (blue) marks nuclei of all cells. INL, inner nuclear layer; ONL, outer nuclear layer; GCL, ganglion cell layer. **(B)** Bipolar cells: Chx10 (red) and PKC-α (white) are markers for bipolar cells; Nrl-GFP (green) and DAPI (blue) as above; INL, ONL, GCL, as above. **(C)** Retinal ganglion cells: Pax6 (white) and Isl1 (magenta) are markers for retinal ganglion cells; DAPI (blue) as above. The white arrows show Isl1 positive retinal ganglion cells in 3D-RA. Calretinin (white) is a marker for inner retinal cells (retinal ganglion cells and amacrine cells); Chx10 (red) and DAPI (blue) as above. Scale, 50 µm for all. **(D) Schematic of anatomy of retinal layers. (E) Three-dimensional in vitro differentiation of iPSC Nrl-*GFP* derived retinal aggregates (3D-RA).** Morphology of iPSCs five days post-thawing (d5). Cells in undifferentiated state are circumferential, "domed shaped" and surrounded by a luminous halo (white arrow). Some unstable colonies tend to differentiate; they have a fibroblastic morphology instead of round shape (red arrow). 3D-RA differentiating retinal aggregates from differentiation DD1 to DD28 obtained following the modified SFEBq protocol [19, 20]. DD1: Rapid re-aggregation of iPSCs after dissociation and passage in a 96-well plate, DD5: Appearance of retinal neuro-epithelium on the retinal aggregates (light edges), DD7: Evagination of retinal neuro-epithelium, DD8-DD20: Growing of evaginating OV like structures, DD22-DD28: GFP expression under control of the *Nrl* promotor in the photoreceptor layer of retinal aggregates. Scale, 400 µm for iPSCs d5; 1000 µm (others). **(F) Dissection of OV-like structures and dissociation of cells into viable single cell solution.** Manual dissection of 3D-RA at stage 2 (DD21). After dissection, the retinal neuro-epithelium tissue was isolated from the pigmented inner cell mass and dissociated into a homogenous solution of single cells. Viability of cells in the solution was 97%. Concentration of cells was 2.0 x 10^6 cells/ml.

### Supplemental figure 2:

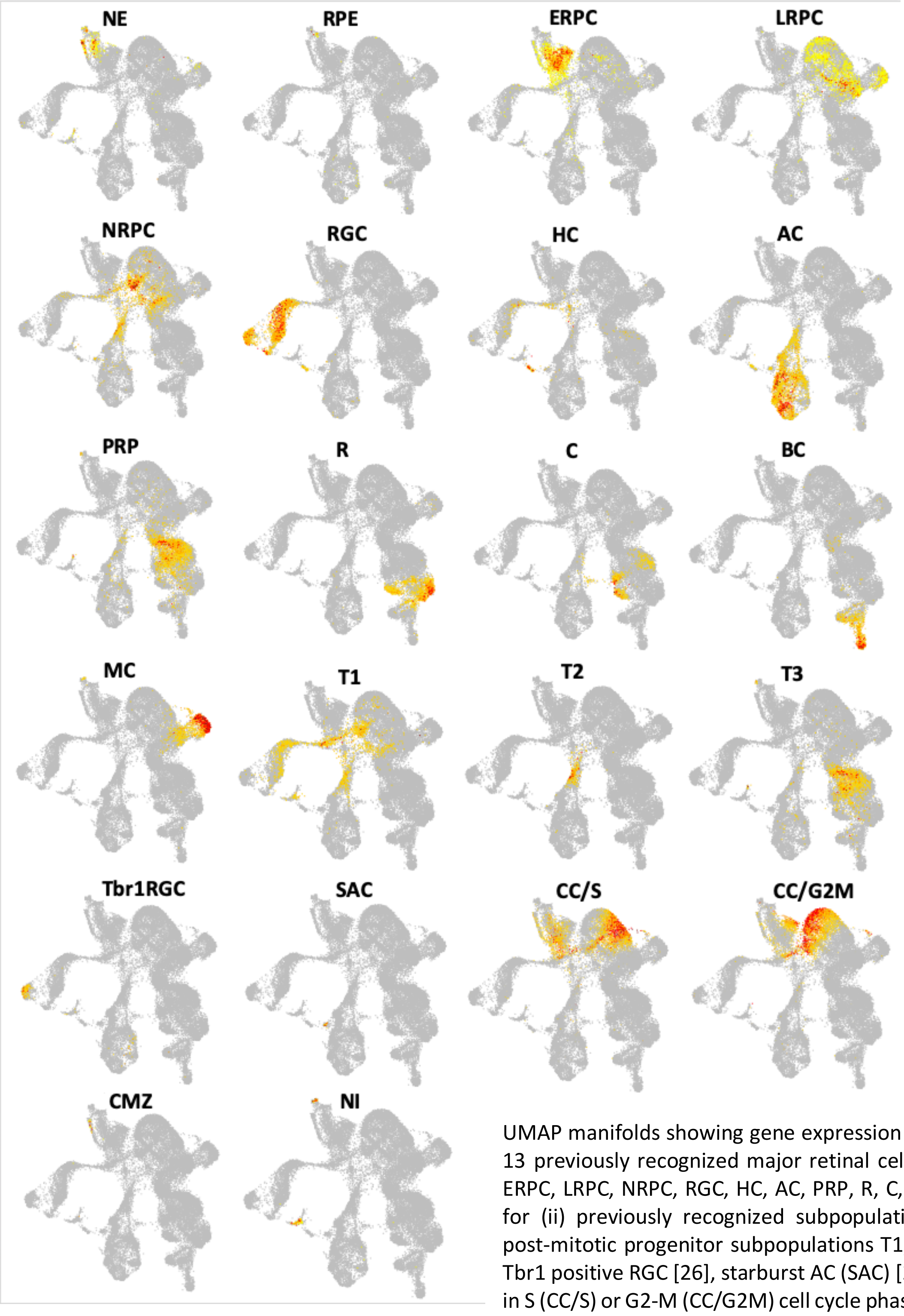

UMAP manifolds showing gene expression signatures for (i) 13 previously recognized major retinal cell types: NE, RPE, ERPC, LRPC, NRPC, RGC, HC, AC, PRP, R, C, BC and MC, and for (ii) previously recognized subpopulations: transitional post-mitotic progenitor subpopulations T1, T2 and T3 [14], Tbr1 positive RGC [26], starburst AC (SAC) [27], dividing cells in S (CC/S) or G2-M (CC/G2M) cell cycle phase [24], the ciliary marginal zone (CMZ) [25], and (iii) a non-identified population (NI) of non-retinal neuronal cells found only in 3D-RA.

### Supplemental figure 3

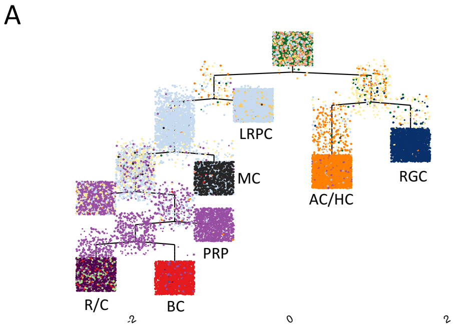

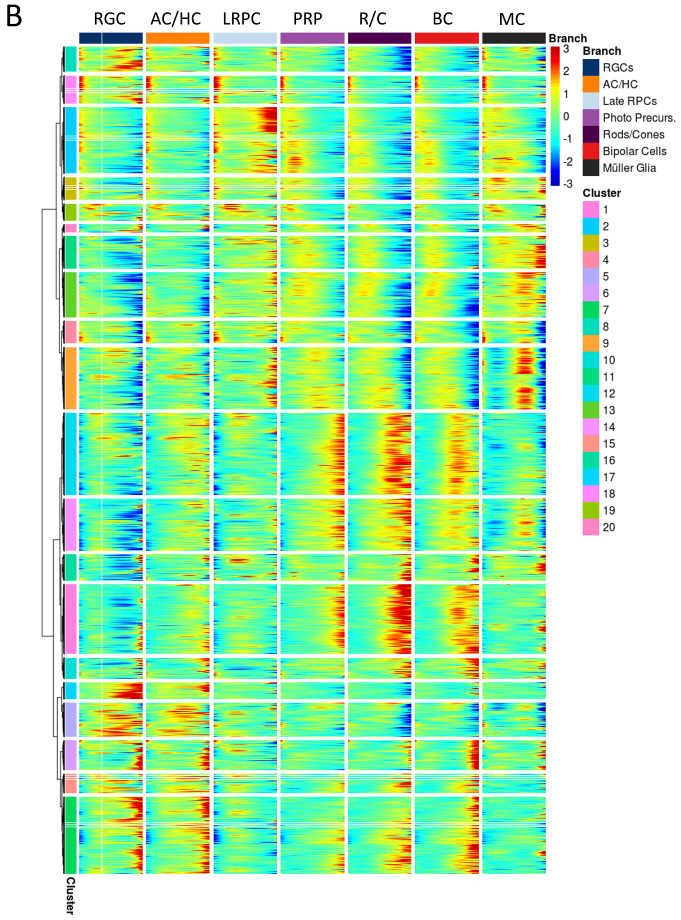

**(A)** Developmental trajectory obtained without supervision with Monocle 2 [12] using NaR cells only. Colors correspond to cell types as defined in the main text and color-coded as in the main figures. As expected, terminal leaves correspond to terminally differentiated cell types (from left to right: R/C, BC, MC, AC/HC and RGC) or to niches of progenitor cells (LRPC, PRP). **(B)** Expression levels (in pseudo-color) along the seven branches of the Monocole 2 tree (X axis) of 10,269 genes that are dynamically regulated as a function of pseudo-time ($q < 0.001$) and grouped in 20 clusters according to their expression pattern (Y axis).

### Supplemental Figure 4:

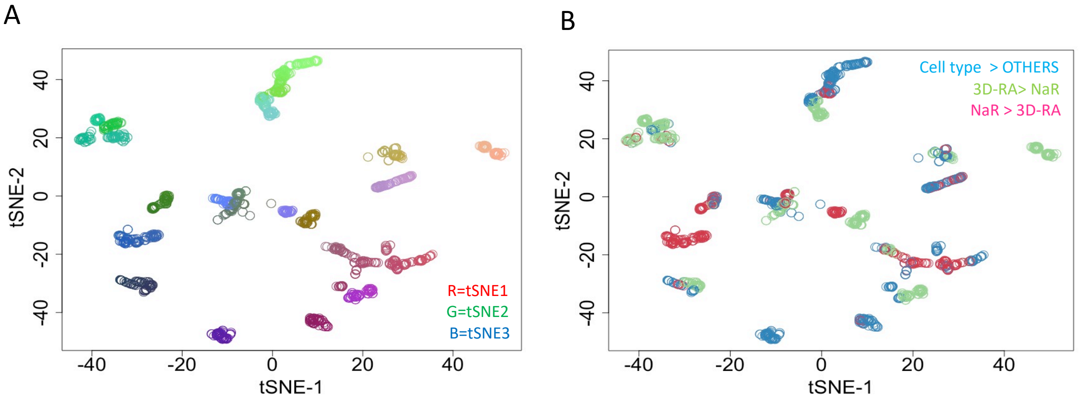

**(A)** tSNE map (dimensions 1 and 2 of 3) of 1,313 gene sets (“found entities”) marking enriched Reactome pathways (each little circle corresponding to a tile in Fig. 3B). Overlapping sets (in terms of gene content) are close to each other in tSNE space (3D). The gene sets are colored in RGB scale where tSNE dimension 1 (tSNE1) determines the intensity of red, tSNE2 the intensity of green, and tSNE3 the intensity of blue. **(B)** Same data points as for (A) but colored by contrast of origin: “Cell type > other”: turquoise; “NaR > 3D-RA”: magenta; “3D-RA > NaR”: lime. One can see that many clusters encompass gene sets corresponding to distinct contrasts, hence highlighting the strong overlap between Reactome pathways that are essential for normal retinal development (“Cell type > other”), and those that are perturbed in 3D-RA relative to NaR (“NaR > 3D-RA” and “3D-RA > NaR”).

### Supplemental figure 5:

Four hundred and twenty-six genes were more strongly expressed in the CMZ (cluster 60 and 69 in Fig. 1A) when compared to all other cell types (STable 4). Strikingly 14 members of the crystallin gene family ranked amongst the top 24 genes (SFig. 5). We noted that the average number of UMIs detected in cells from cluster 60 was nearly twice the experiment-wide average (p = 0.01). We therefore surmise that this cluster contains a large proportion of doublets of physically connected neuro-retinal and posterior epithelial lens cells, explaining the detection of crystallin transcripts and the marked separation of clusters 60 and 69 from the rest of NE and ERPC.

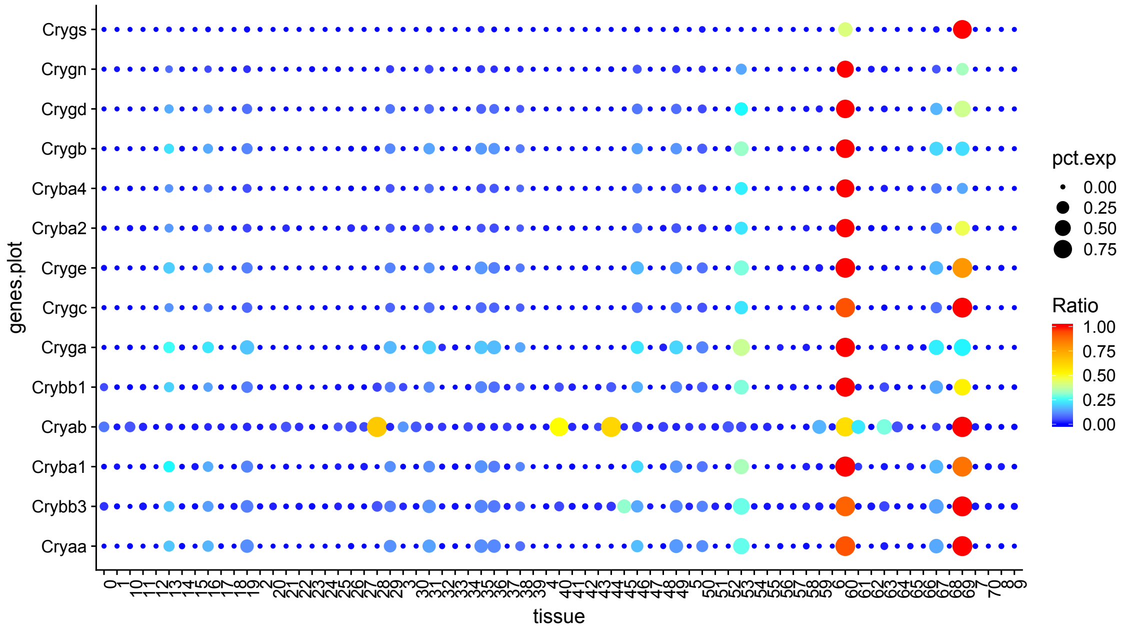

Level of expression in all 70 clusters of 14 crystallin genes. The 14 chosen crystallin genes are the ones ranked amongst the top 24 genes differentially expressed in the CMZ (=cluster 60 + cluster 69) compared to all other cell types.

### Supplemental Figure 6:

A

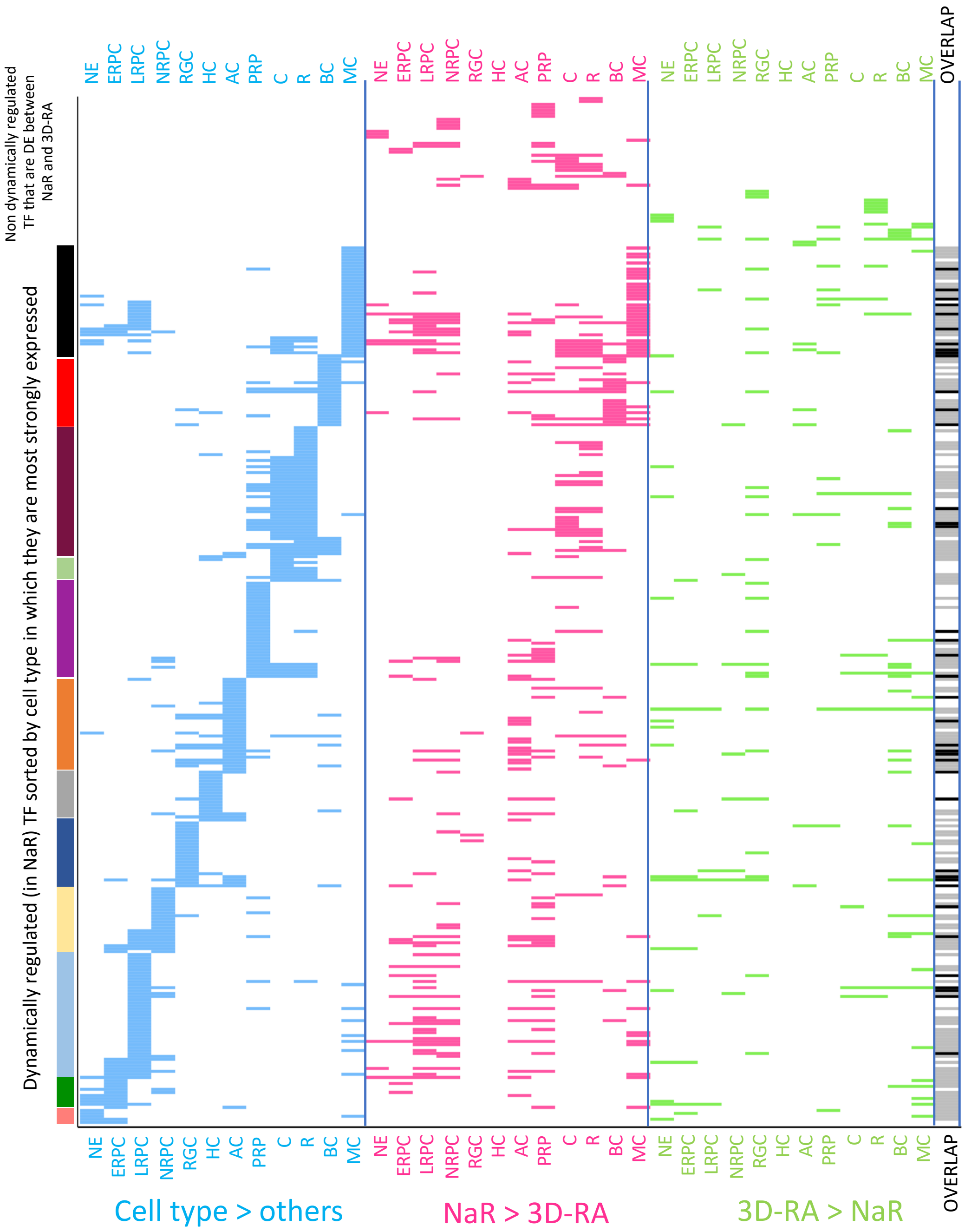

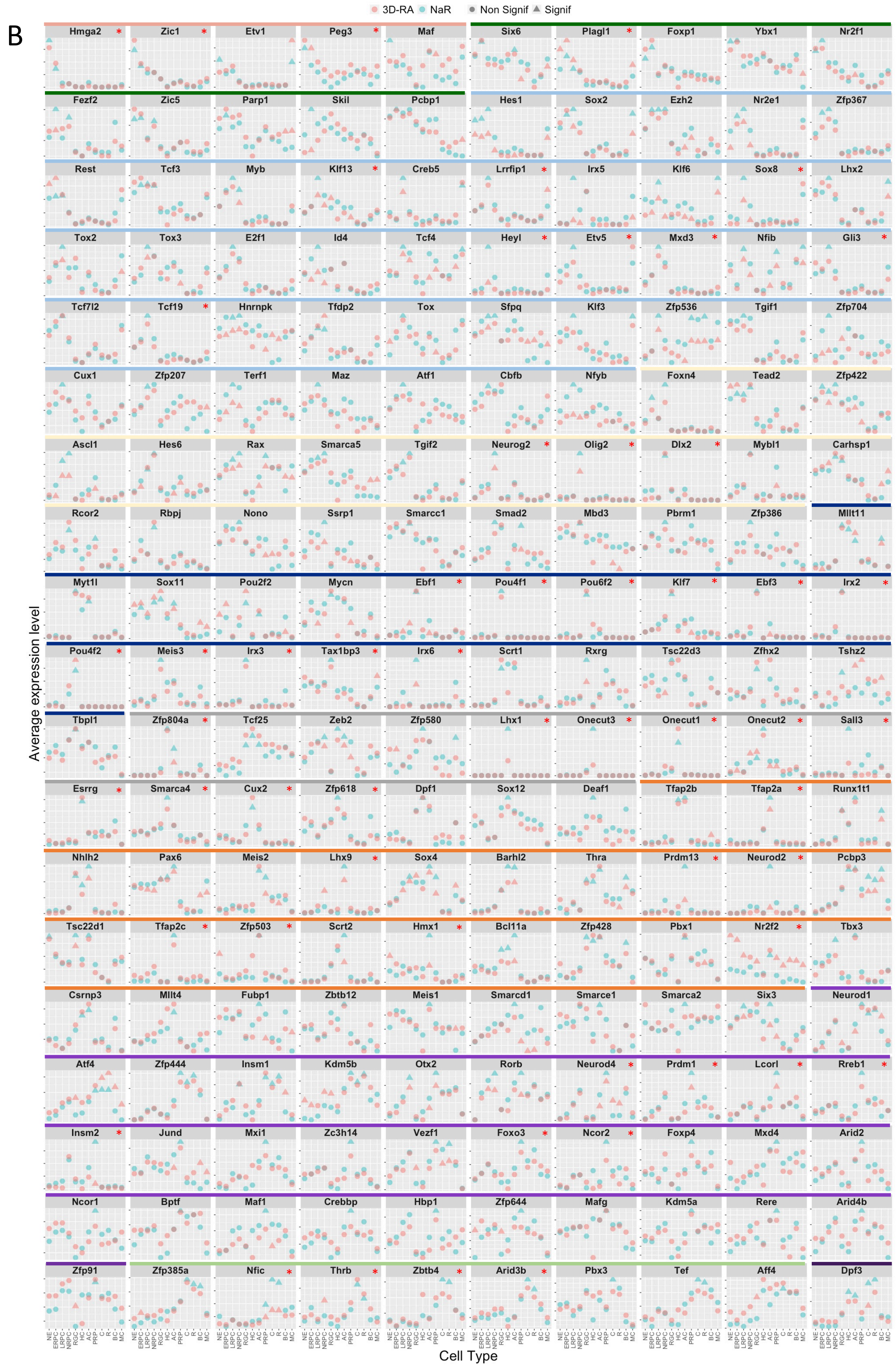
B
3D-RA
NaR
Non Signif
Signif
Average expression level
Cell Type
NE
ERPC
LRPC
NRPC
RGC
HC
AC
PRP
C
R
BC
MC
Hmga2
Zic1
Etv1
Peg3
Maf
Six6
Plagl1
Foxp1
Ybx1
Nr2f1
Fezf2
Zic5
Parp1
Skil
Pcbp1
Hes1
Sox2
Ezh2
Nr2e1
Zfp367
Rest
Tcf3
Myb
Klf13
Creb5
Lrrfip1
Irx5
Klf6
Sox8
Lhx2
Tox2
Tox3
E2f1
Id4
Tcf4
Heyl
Etv5
Mxd3
Nfib
Gli3
Tcf7l2
Tcf19
Hnrnpk
Tfdp2
Tox
Sfpq
Klf3
Zfp536
Tgif1
Zfp704
Cux1
Zfp207
Terf1
Maz
Atf1
Cbfb
Nfyb
Foxn4
Tead2
Zfp422
Ascl1
Hes6
Rax
Smarca5
Tgif2
Neurog2
Olig2
Dlx2
Mybl1
Carhsp1
Rcor2
Rbpj
Nono
Ssrp1
Smarcc1
Smad2
Mbd3
Pbrm1
Zfp386
Mllt11
Myt1l
Sox11
Pou2f2
Mycn
Ebf1
Pou4f1
Pou6f2
Klf7
Ebf3
Irx2
Pou4f2
Meis3
Irx3
Tax1bp3
Irx6
Scrt1
Rxrg
Tsc22d3
Zfhx2
Tshz2
Tbpl1
Zfp804a
Tcf25
Zeb2
Zfp580
Lhx1
Onecut3
Onecut1
Onecut2
Sall3
Esrrg
Smarca4
Cux2
Zfp618
Dpf1
Sox12
Deaf1
Tfap2b
Tfap2a
Runx1t1
Nhlh2
Pax6
Meis2
Lhx9
Sox4
Barhl2
Thra
Prdm13
Neurod2
Pcbp3
Tsc22d1
Tfap2c
Zfp503
Scrt2
Hmx1
Bcl11a
Zfp428
Pbx1
Nr2f2
Tbx3
Csrnp3
Mllt4
Fubp1
Zbtb12
Meis1
Smarcd1
Smarce1
Smarca2
Six3
Neurod1
Atf4
Zfp444
Insm1
Kdm5b
Otx2
Rorb
Neurod4
Prdm1
Lcorl
Rreb1
Insm2
Jund
Mxi1
Zc3h14
Vezf1
Foxo3
Ncor2
Foxp4
Mxd4
Arid2
Ncor1
Bptf
Maf1
Crebbp
Hbp1
Zfp644
Mafg
Kdm5a
Rere
Arid4b
Zfp91
Zfp385a
Nfic
Thrb
Zbtb4
Arid3b
Pbx3
Tef
Aff4
Dpf3

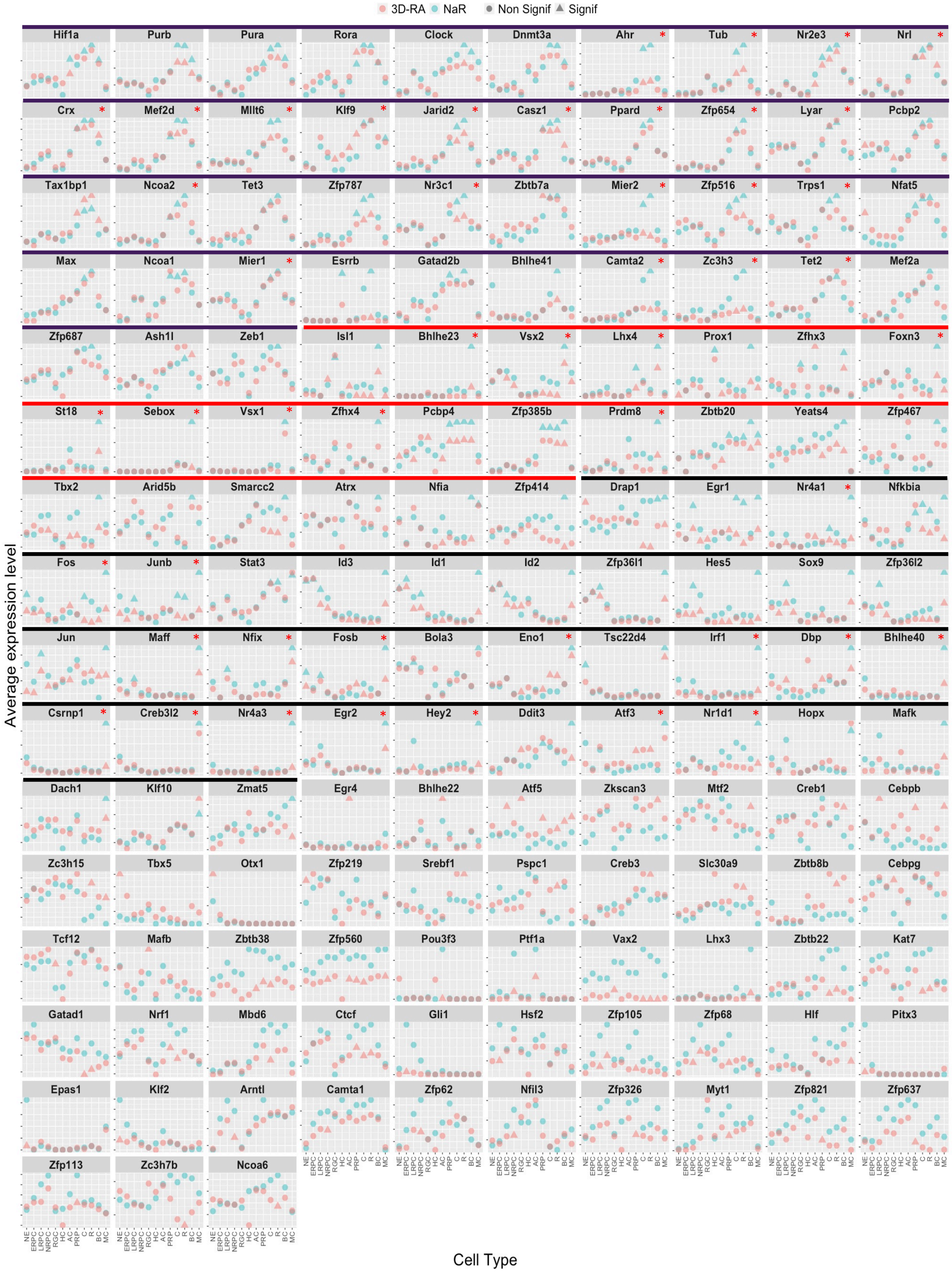

**(A)** Transcription factors (TF) that are (i) differentially regulated between cell types (turquoise), (ii) under-expressed in 3D-RA when compared to NaR (magenta), or (iii) over-expressed in 3D-RA when compared to NaR (lime). OVERLAP: TF that are differentially expressed in the three conditions (Cell type > others, NaR>3D-RA and 3D-RA> NaR) are marked in black. TF that are differentially expressed during retinal development (Cell type > others) and in one of the NaR vs 3D-RA conditions (NaR>3D-RA or 3D-RA> NaR) are marked in grey. Acronyms for cell types are as in the remainder of the manuscript. **(B)** Average expression level (% UMI) in 12 retinal cell types in NaR (green) and 3D-RA (red) of the 343 TF (cfr. A) that are significantly overexpressed in at least one cell type when compared to all others in NaR (293) and/or significantly over- or under-expressed in 3D-RA when compared to NaR in at least one cell type. Green triangles mark cell types in which the corresponding gene is significantly (q < 0.01, i.e. accounting for multiple testing) overexpressed in NaR when compared to all other cell types combined. Red triangles mark cell types in which the expression level differs significantly (q < 0.01) between Nar and 3D-RA. The gene names are given in the facet headers. The cell type in which the TF is overexpressed (NaR) are indicated by the colored bars (color code as in A and other figures). Red asterisks mark the genes that were selected for Fig. 4A.

### Supplemental Figure 7:

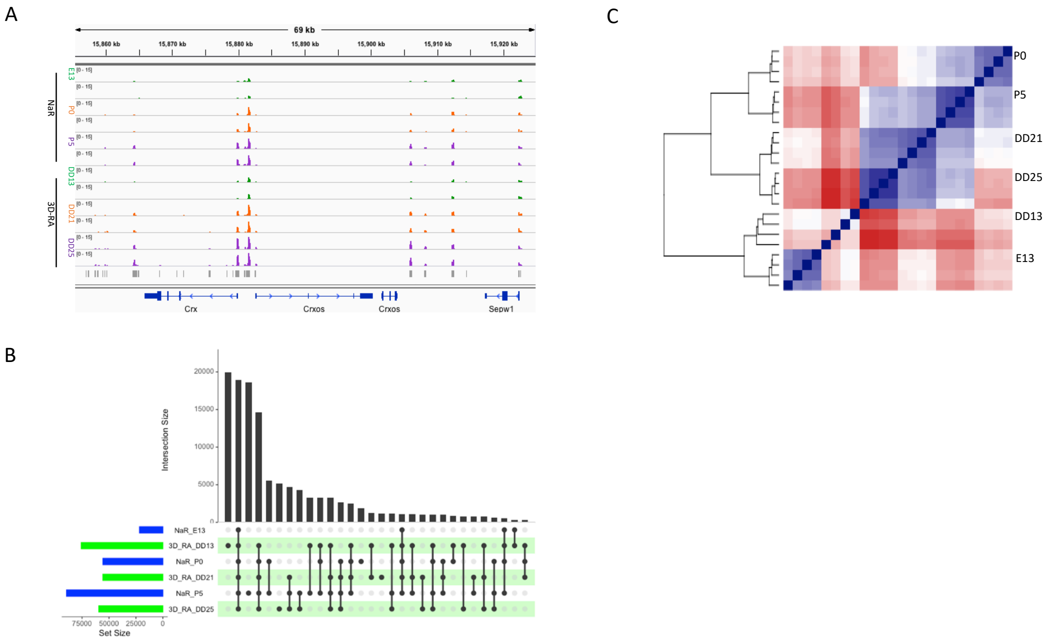

**(A) Example of ATAC-Seq results obtained in NaR and 3D-RA samples in the vicinity of the *Crx* gene.** The intensity of several peaks is increasing with developmental stage in both NaR and 3D-RA as expected for this TF that is primarily expressed in PRP, C and R. **(B) Numbers and overlap between ATAC-Seq peaks detected in different sample types**. The number of peaks increases with developmental stage in NaR but not in 3D-RA. A large proportion of peaks are either DD13- (16.1%) or P5-specific (15.1%), or shared between all samples (15.3%), or all samples minus E13 (11.9%). **(C) Hierarchical clustering of the samples based on the intensity of 123,482 ATAC-Seq peaks.** Biological and technical replicates always cluster together by sets of four. Sample types cluster by developmental stage at stage I (E13 and D13), but by origin (NaR vs 3D-RA) at stages II and II. Clustering was based on Pearson's correlation coefficient and used Ward.D's method. The samples are in the same order on the X and Y axis and reflecting according to the resultingdendrogram

### Supplemental Figure 8:

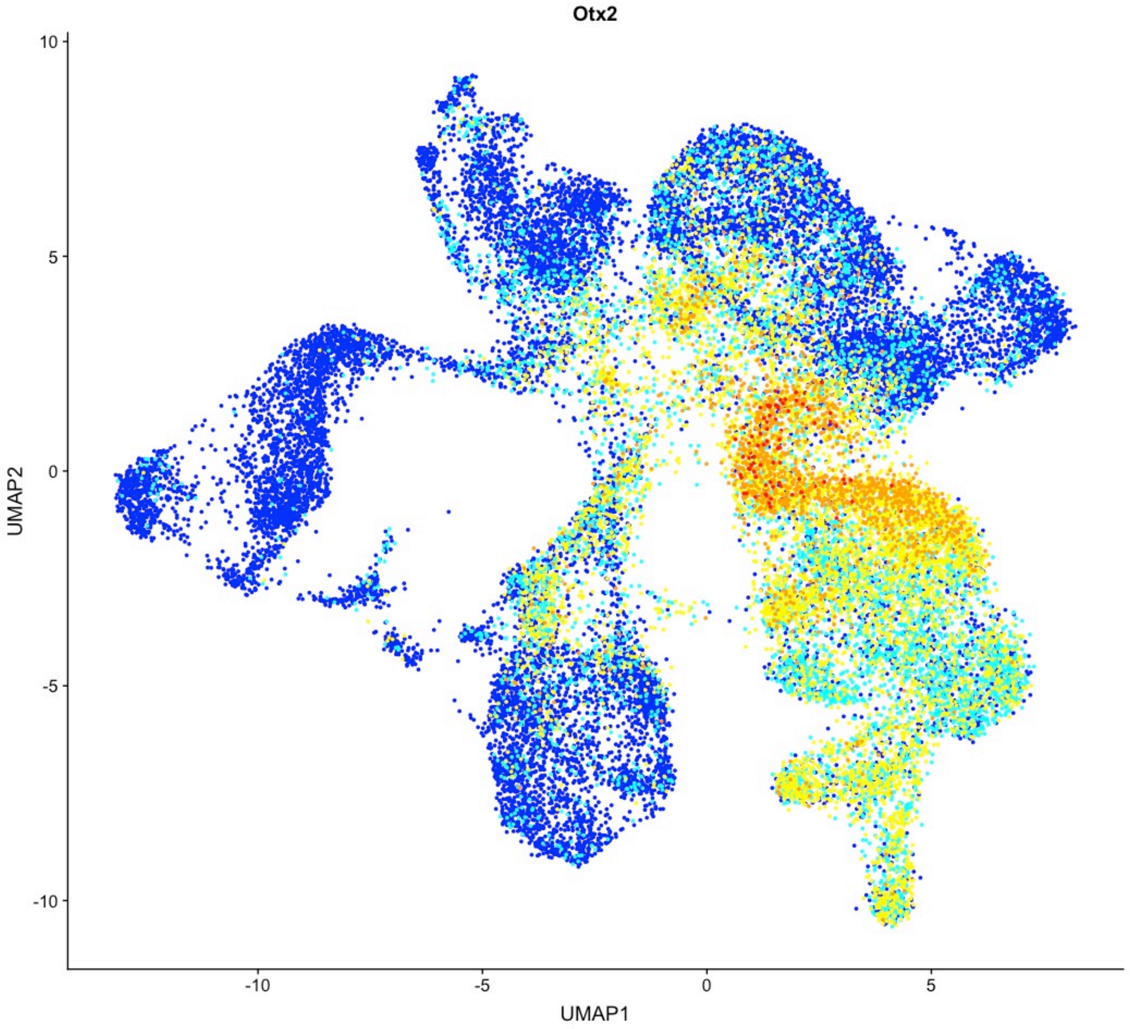

Illustration of the fact that the NRPC cells that express Otx2 are primarily concentrated in the vicinity of PRP cells.

### Supplemental Figure 9:

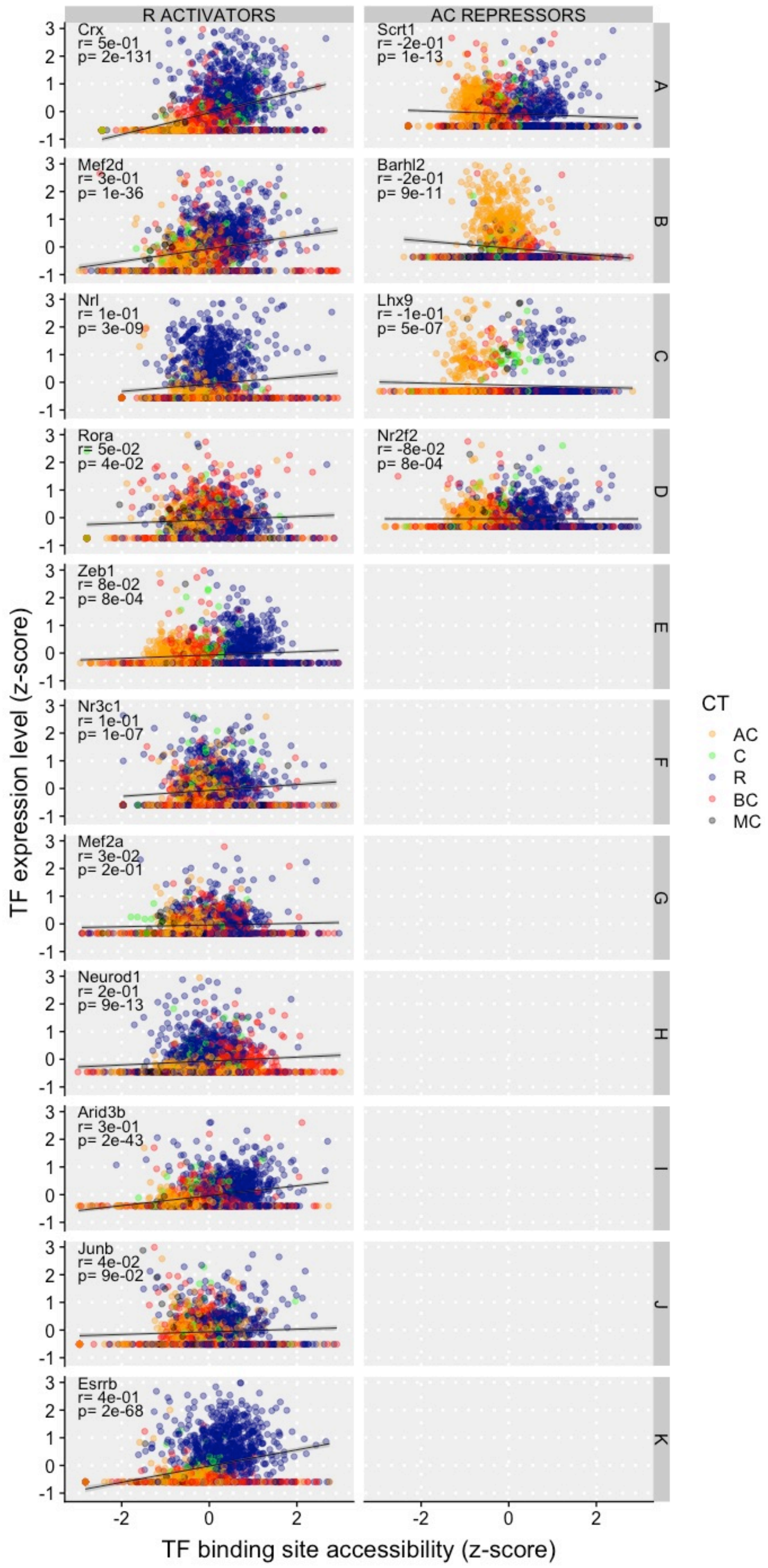

Scatter plots showing the correlation between gene-specific TF transposase accessibility (y axis) and corresponding genome-wide TF binding motif transposase accessibility (x axis) 11 presumed R activators and 4 AC repressors. All correlations are positive for the R activators, and negative for the AC repressors, and significant for 13/15.

### Supplemental Figure 10:

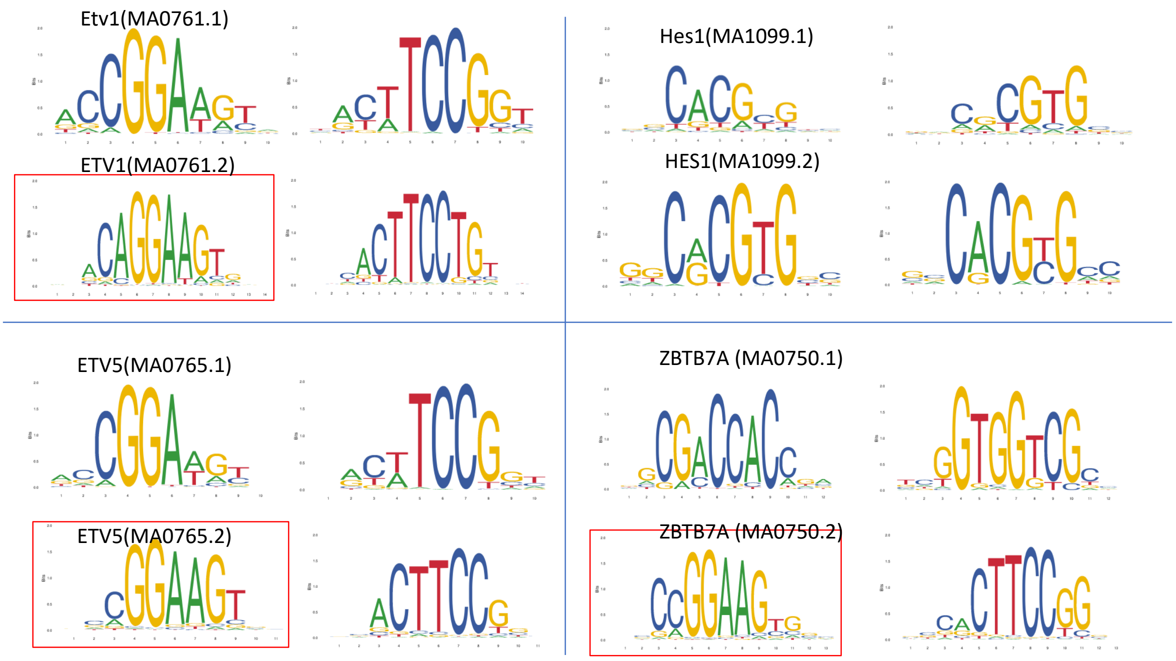

Binding motifs for Etv1, Etv5, HJes1 and Zbtb7a.

### Supplemental Figure 11:

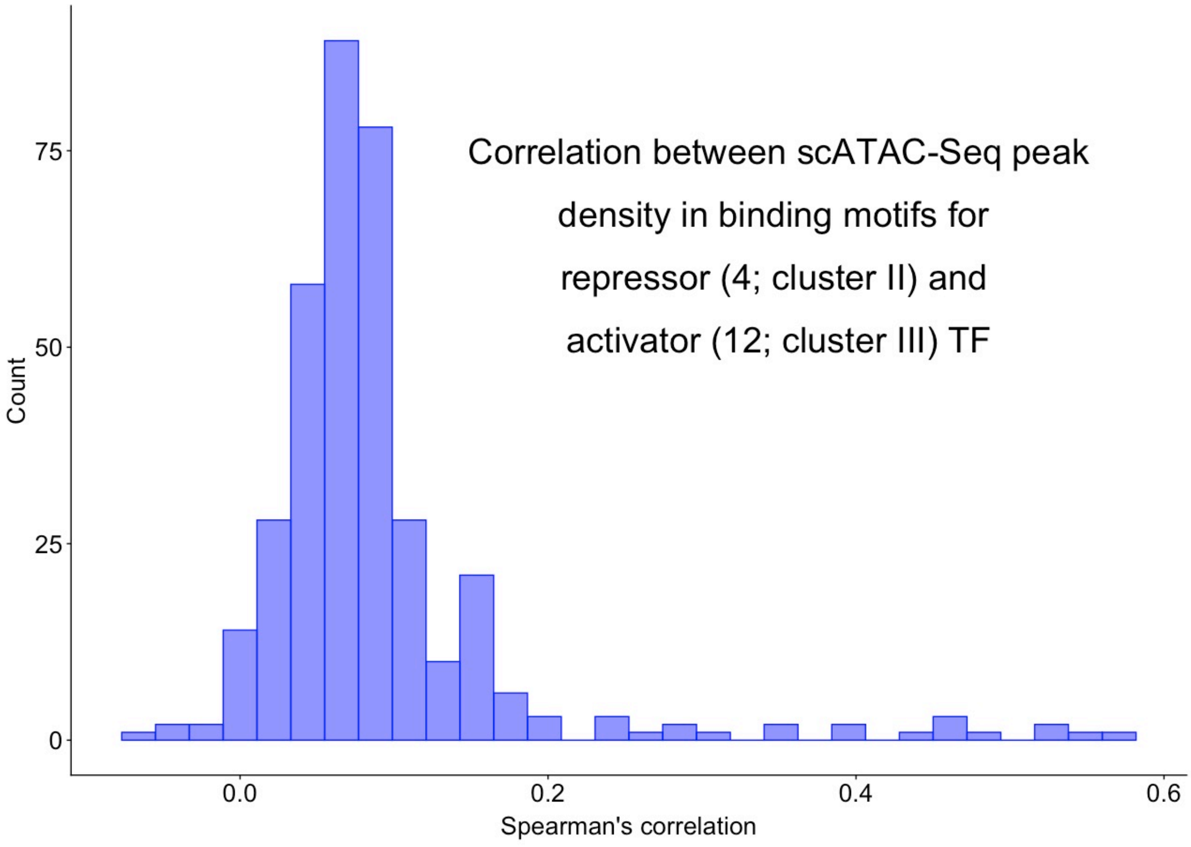

Pair-wise correlations (Spearman) between the density (number of binding motifs divided by peak size) in binding motifs in scATAC-Seq peaks [39] for 4 repressor TF active in AC (cluster II)( Barhl2, Lhx9, Nr2f2, Scrt1) and 11 activator TF active in C and R (cluster III)( Arid3b, Crx, Esrrb, Junb, Mef2a, Mef2d, Neurod1, Nrl, Nr3c1, Rora, Zeb1).
